# Melanoma innervation is associated with cancer progression in a zebrafish xenograft model

**DOI:** 10.1101/2023.12.13.571512

**Authors:** Francesca Lorenzini, Johanna Marines, Julien Le Friec, Nam Do Khoa, Maria Angela Nieto, Berta Sanchez-Laorden, Maria Caterina Mione, Laura Fontenille, Karima Kissa

## Abstract

The peripheral nervous system has a key role in regulating tumour biology in different types of cancer. Here, by modelling aggressive melanoma in larval zebrafish xenografts, we highlight the dynamics of tumour innervation in the tumour microenvironment (TME). Axonogenesis and dendritogenesis are detected in the motoneurons surrounding the melanoma niche and neurogenesis is observed in the nearby population of the enteric nervous system. We also demonstrate the crucial role of catecholamines in promoting melanoma progression, supporting *in vivo* cancer cell dissemination and invasion. This *zebrafish* model will allow to uncover neural markers associated with melanoma progression to help in the design of innovative anti-neurogenic therapies targeting specifically the neuronal signals that regulate melanoma progression.

**Graphical abstract:** 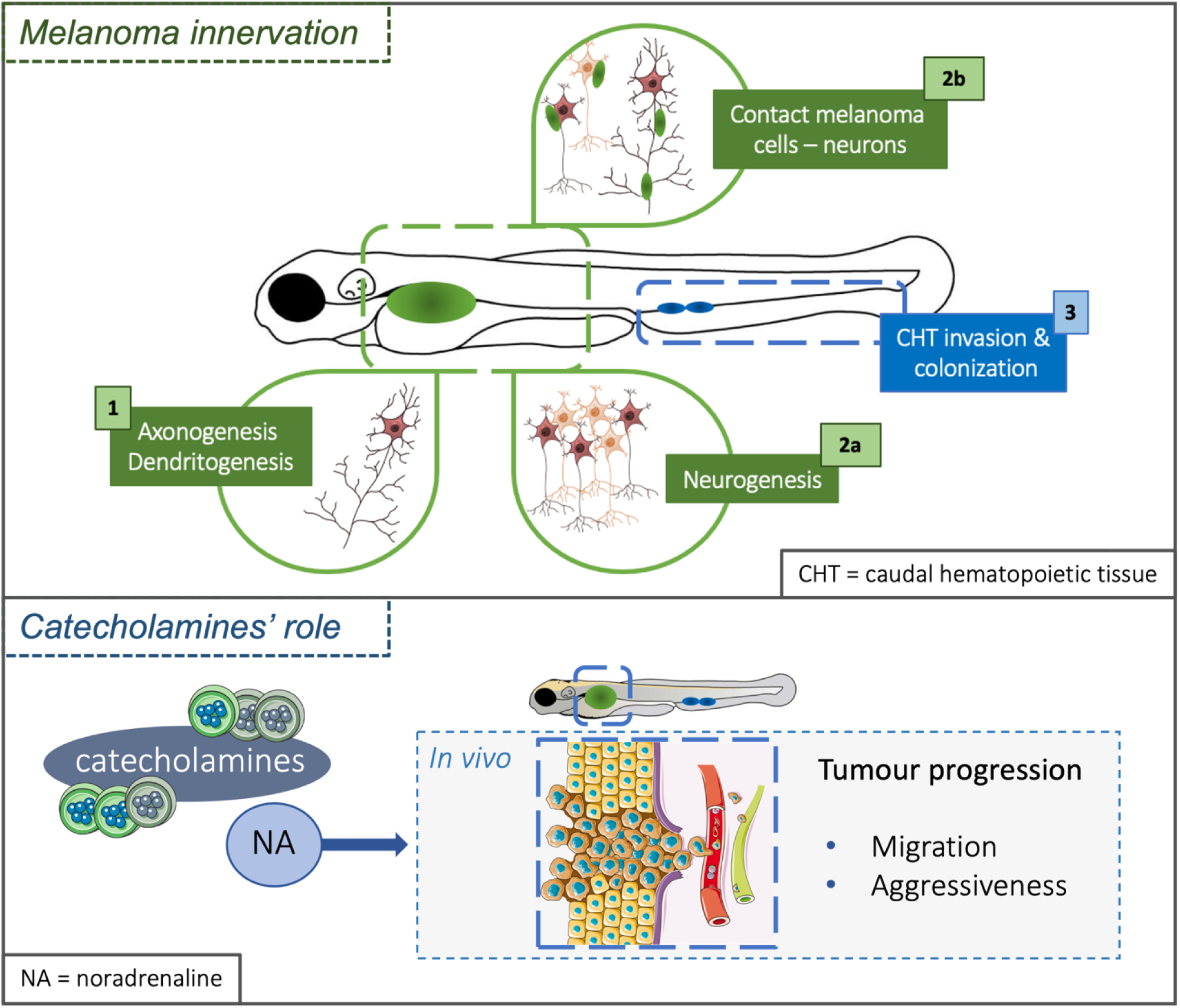

**Highlights:**

- Transplantation of human melanoma cells in 3 dpf zebrafish swim bladder allows the development of aggressive melanoma, which cells invade the surrounding organs and migrate over distant locations.
- The presence of melanoma cells in the larval zebrafish induces morphological changes in the motoneurons inside the tumour niche, including increased axon length and dendritic arborization.
- The invasion of melanoma cells in the larval intestine promotes neurogenesis of enteric neurons.
- Transplanted melanoma cells display direct contact with enteric neurons in the intestinal region and migrate along axons to escape from the primary cancer mass, as a mechanism similar to vessel co-option during metastatic dissemination.
- Catecholamines promote melanoma cell migration and invasion in the zebrafish, modelling melanoma progression.

## Introduction

Cutaneous melanoma is a cancer that arises from skin melanocytes, the pigment-producing cells deriving from neural crest cells. It is the most aggressive type of skin cancer, due to its high metastatic capacity. Indeed, although it represents only 1% of all skin cancers, it causes the majority of the deaths related to skin cancers ^1^. In the last 15 years, the development of targeted and immunotherapies have improved the survival of metastatic melanoma patients ^2,3^. However, most patients develop resistance to therapies, due to the high plasticity of melanoma cells ^4,5^ and the presence of a very dynamic tumour microenvironment (TME).

Among the non-cancer cells that populate the melanoma TME, the functions of macrophages, vascular endothelial cells and fibroblasts have been extensively studied ^6,7,8^. However, the role of a novel player, the peripheral nervous system (PNS) in the melanoma TME is poorly understood. The PNS has been recently associated with carcinogenesis and cancer progression, through the release of different factors in several types of tumours. Moreover, cancer cells can physically use the nerve fibres as route of cancer dissemination, a mechanism that is called perineural invasion (PNI) ^9,10^. Simultaneously, cancer cells can also impact the neural network in the TME, and all these modifications are due to the plasticity of the NS. The changes are induced by the activation of neurogenesis and axonogenesis^11^, mechanisms that are fundamental during embryonic development. Neurons can also communicate with other cells inside the tumour niche, shaping a pro-tumoral TME by inducing immunosuppression and/or angiogenesis ^12,13^.

The autonomic branch of the peripheral nervous system (PNS), composed of the sympathetic nervous system, the parasympathetic nervous system, and the enteric nervous system, is frequently associated with solid cancers ^14,15,16,17,18^. As such, the sympathetic and parasympathetic NS have been respectively associated with prostate cancer initiation and progression ^14^. Tumour innervation has been also reported for other cancer types, including gastric, pancreatic, skin, breast, colon and ovarian carcinomas ^18^. However, the contribution of innervation to melanoma is not that well characterized. It has been shown that sensory neurons counteract melanoma progression ^19, 20^. However, nociceptor neurons, present in melanoma samples from patients, contribute to melanoma growth in mice by decreasing antitumour immune responses ^21^. In addition, some preclinical studies have demonstrated the role of autonomic NS neurotransmitters such as adrenaline, noradrenaline, aka catecholamines, and acetylcholine in cancer progression ^22,23^. Indeed, β1, β2 and β3 catecholamine receptors, physiologically expressed in different tissues and organs, to regulate vital body functions have been also detected in the membrane human melanoma samples and melanoma cell lines ^24,25,26^. Activation of β-receptors seem to support cell proliferation, while inducing migration and invasion phenotypes and an EMT-like process in melanoma cells *in vitro* ^24,27^. Recently, a clinicopathological study demonstrated that patients developing a melanoma with high level of β2-receptor expression presented more aggressive and advanced stages of the disease. In addition, β-blockers, antagonists of β-adrenergic receptors, have positive, anticancer effects in the treatment of melanoma patients ^28^. All of this supports an active role of adrenergic signalling in melanoma progression. Therefore studying the relevance of β-adrenergic signaling and tumour innervation, both as a diagnostic and prognostic factor, as well as in the perspective of developing innovative anti-neurogenic therapies is of utmost importance. However, tumour innervation in melanoma has never been imaged in *in vivo* pre-clinical models.

Xenografts in larval zebrafish have emerged as a very powerful tool to characterize the role of different components of the TME ^29^, thanks to the availability of many transgenic lines bearing reporters of different cells in the TME including neurons. Here we are using a human melanoma xenograft model in the zebrafish embryo that we have recently developed ^30^. We report that this model recapitulates melanoma progression to the invasive state. Melanoma cells were able to escape from the primary mass and disseminate to near and distant locations. Using real time imaging, we have investigated the presence of tumour innervation and revealed the dynamics of tumour-induced neurogenesis and axonogenesis, and the *in vivo* pro-tumoral role of catecholamines. Our *in vivo* model will allow testing new drugs, including β-blockers, in the early, medium, and late stages of cutaneous melanoma.

## Materials and methods

### Chemical compound used both for *in vivo* and *in vitro* experiments

Epinephrine Bitartrate (AD) and L-(−)-Norepinephrine (+)-bitartrate salt monohydrate (NA) were purchased from Merck. Before all experimental procedures fresh drug’s solutions were prepared by diluting the compounds in purified water. Zebrafish larvae were transplanted with A375P cells dissolved in PBS with either 10 uM NA or 1 uM AD. Control larvae (CTL) were transplanted with A375P in PBS.

### Animal rearing

Zebrafish (*Danio rerio*) strains were raised and maintained in the Fish Facility of AZELEAD under standard conditions ^31^. Adult zebrafish were maintained on a 12/12 h light/dark cycle in a partially recirculating system and fed 3 times/day with fresh Artemia salina and dry food. Transgenic lines (***Table 1***) and wildtype strains were crossed to obtain offspring for experimental procedures. Incrosses of wildtype strains and homozygous transgenic line *tg(NBT:dsRed)* were done to have offspring for cell transplantation procedure. Adult *Et(kita:GalTA4,UAS:mCherry)^hzm1^*; *Tg(UAS:eGFP-HRAS^GV12^)^io6^* zebrafish were raised to generate adult zebrafish with melanoma tumours. All experimental procedures on zebrafish were performed in Fish Facility in accordance with the European guidelines and regulations on Animal Protection from the French Ministry of Health (F341725).

### Cell culture maintenance

Human primary melanoma cell line, A375P, was cultured in Dulbecco’s modified Eagle’s medium (DMEM) (Eurobio scientific) supplemented with 1% L-glutamine 200mM (Eurobio scientific), 1% Penicillin/Streptomycin (Eurobioscientific #CABPES01-0U) and 10% foetal bovine serum (FBS) (Eurobio scientific) at standard conditions of 5% CO2, at 37°C. Cells were cultured in 100 mm Petri dish (Corning) and split twice a week when they reached 80-90% confluency.

### Generation of stable fluorescent human cancer cell lines

To obtain A375P-eGFP+ cells, A375P cells were stably transfected with sfGFP-N1 as described below. sfGFP-N1 (Addgene plasmid #54737) plasmid was a gift from Geoffrey Waldo. One day before transfection, cells were plated in 6 well plates. sfGFP-N1 purified plasmid (4 μg) was transfected using 2.5 M CaCl2 reagent. 24h after transfection, complete DMEM medium was replaced. eGFP+ cells were selected using Geneticin (800-1000μg/mL) diluted in DMEM medium for 4-6 weeks before starting *in vivo* experiments.

### Embryo and cell preparation prior to xenotransplantation

Embryos were initially maintained at 28 °C at maximum density of 50 embryos per Petri dish in fish water supplemented with 0.0002% methylene blue (Sigma) as an antifungal agent. After 24h, embryos were placed in fish water containing 200 μM Phenylthiourea (PTU) to prevent embryo pigmentation and allow fluorescent imaging acquisition.

Fluorescent human melanoma A375P-eGFP+ cells were harvested from a 100 mm Petri dish the day of xenograft transplantation. Cells were washed with phosphate-buffered saline (PBS) and detached using Trypsine - Versene EDTA (Eurobio scientific) at 37°C for 5 min. Trypsine - Versene was inactivated by complete DMEM medium, and cells were centrifuged at 1000 rpm for 5 min and then washed with PBS. In some experiments, to study the *in vivo* role of catecholamines on melanoma progression, catecholamines (either 10 uM NA or 1 uM AD) were added in the PBS. Finally, cells were resuspended in 50-100 μL PBS to ensure highly concentrated cell preparation.

### Xenotransplantation of human cancer cell lines

Borosilicate glass capillaries (O.D.: 1 mm, I.D.: 0,75 mm, Sutter Instrument) were pulled using glass micropipette puller (Sutter Instrument). Capillaries were filled with 10 μL of cells suspension. Injection was performed under a stereomicroscope (Leica M80 Stereo zoom microscope) using a micromanipulator (Narishige) and the oil manual microinjector (Cell Tram Vario Eppendorf). Before xenografting, 3 dpf larvae were anesthetized in 0.16 mg/ml PBS/Tricaine (MS-222). 200 to 400 melanoma cells were transplanted in the nascent swim bladder. Control larvae were injected with PBS in the same region. Larvae with fluorescent tumour mass in the swim bladder were selected and isolated in a single well of 24-well plates. Transplanted embryos were maintained at 33°C in fish water containing PTU.

### Live-imaging acquisition

Transplanted larvae were individualized and imaged the day of transplantation (D0), two days after transplantation (D2), and either 3 (D3) or 4 days post transplantation (D4), according to the biological question of the experimental procedure. D0 corresponds to the day of transplantation with a technical delay of about 6h between the transplantation procedures and the imaging acquisition. For image acquisition, every larva was anaesthetized and positioned laterally inside a single well of a 7-position mould (designed by AZELEAD) covered with fish water containing Tricaine. To study tumour progression and tumour innervation, live images were acquired with LSM 510 Meta Zeiss confocal microscope (Objective: 10X/1). 488 and 543 nm lasers were used with appropriate filters. Images were taken with 5,7 μm z-step interval to a total Z-stack of 170-200 μm depending on tumour size.

To study the effect of catecholamines on tumour growth and tumour invasiveness, images were acquired using the Zeiss automated microscope system Celldiscoverer 7 (CD7) (Objetives: 5x/1 and 20x/0,5). Images were acquired in z-stack mode inside a total range of 250 μm with a z-step interval of 5 μm. 470 nm LED was used to acquire green fluorescence.

### Image analysis

Images were processed using Fiji and NeuronStudio software. Maximum projections were generated using either Fiji software or ZEN software for the images acquired using the CD7 microscope.

Melanoma growth and evolution were followed thanks to green fluorescence of A375P melanoma cancer cells. To analyse tumour growth, the maximum projection of every image was used. To determine tumour area, a threshold was applied in Fiji for each larva on D0, D2 and D3/D4 to generate a mask. The tumour area was then normalized on the day of transplantation (D0) to assess the percentage of tumour growth.

To characterize the migratory and invasive potential of melanoma cancer cells *in vivo*, disseminated cancer cells from the primary mass and elongated cells were manually counted on Fiji moving through the entire Z-stack. To check the presence of cancer cells in the caudal hematopoietic tissue (CHT), A375P cells were manually counted from the urogenital opening till the posterior part of the CHT.

To study tumour innervation, NeuronStudio software was used to manually trace in 3D five selected axons, named from 1 to 5, that located all around the region of swim bladder.

The length of major axons and the number of branching points were measured and counted. To describe the dendritic anatomy, Sholl analysis was performed. Sholl analysis counts the number of the intersections of the dendrites with different imaginary rings, reporting the distance from the soma of the neurons, considered as the centre. The number of intersections is the number of the dendritic branches. The data acquired from the analysis allow the construction of a graph called Sholl profile.

The images were prepared using the Fiji software by adjusting brightness and contrast.

### Zebrafish biopsies

Adult wild type and *kita:ras* fish were sacrificed by anaesthetic overdose (0,32 mg/ml MS-222) and immediately put on a homemade sectioning stage containing dry ice under a stereomicroscope (Leica M80 Stereo zoom microscope). Using scalpel and forceps, dark melanoma biopsies from *kita:ras* fish were dissected. Melanoma came from either the central body or the tail of adult fish. As control, biopsies from wild type fish were obtained in the same way. Melanoma and wild type biopsies were immediately snap frozen using dry ice after dissection. Samples were stored in 2 ml Eppendorf in liquid nitrogen.

### RNA extraction, cDNA synthesis and qPCR – zebrafish samples

Transplanted larvae (MT) with A375P cells and control larvae (CTL) injected with PBS on D0 and D4 were anesthetized (1x MS-222). Then larvae were beheaded under the stereomicroscope with a sterile scalpel. 10 beheaded larvae were grouped for every condition (CTL D0, MT D0, CTL D4 and MT D4) in 2 ml Eppendorf and snap frozen in dry ice. The samples were then stored at -80 °C until RNA extraction procedure.

Same RNA extraction was done on larvae samples and on adult zebrafish biopsies. RNA extraction was performed using NucleoSpin RNA purification kit (MACHEREI-NAGEL), according to the manufacturer’s protocol. At the final step, RNA was eluted in 30 μl RNase-free water to increase its concentration. Quantity and quality of RNA were assessed through NanoDrop 2000c (Thermo Scientific). Each sample was stored at -80 °C. 400 ng of total RNA was retrotranscribed to cDNA using SensiFAST^TM^ cDNA Synthesis Kit (Bioline), according to the manufacturer’s protocol. Quantitative PCR reaction mix was prepared using 2x Fast Q-PCR Master Mix (SYBR, no ROX, SMOBIO), according to the manufacturer’s protocol. cDNA was diluted 1:4 with nuclease-free water. qPCR reactions were performed on a CFX96 Real-Time PCR Detection System (Bio-Rad) machine to amplify the cDNA of human and zebrafish genes of interest. Primers were designed with Primer 3 program and were purchased from Eurofins Genomics; the sequences are reported in ***Table 2***. The primers’ specificity was tested checking the melting curves. The analysis of gene expression is conducted using the ΔΔCt mathematical methodology on Microsoft Excel program.

### Statistical analysis

Statistical analyses were conducted using GraphPad Prism 8 software. Results were shown as mean ± SEM (Standard Error of the Mean), percentage mean ± SEM and in violin plots (centre lines, medians; box limits, second and third quartiles). Differences among groups were analysed by two-tailed unpaired/paired Student’s t test, non-parametric test, One-way/Two-way ANOVA and Friedman test, as indicated in the figures’ legends. According to the statistical test, significance was set at either P-value <0,05 or P-value <0,0332, as indicated in the figures’ legend.

## Results

### Melanoma xenografts in larval zebrafish recapitulates steps of an aggressive cancer

To evaluate the presence and role of innervation in melanoma progression, we used a novel transplantation assay, developed in zebrafish and described in *Lorenzini et al*, 2023 ^30^. Green fluorescent human malignant melanoma cell line (A375P) were transplanted in the swim bladder (SB) of 3 dpf larvae (*Figure 1A*). Live-image acquisition was performed from the day of transplantation (D0) up to 4 days post transplantation (D4) using either a fluorescent microscope or a confocal microscope (*Figure 1B*). Melanoma cells were able to proliferate, establishing a solid tumour mass that increased its size over time (*Figure 1B, 1C;* n=20/21). Significant melanoma growth was observed from D0 to D2 (** P-value < 0,0061) and also from D0 to D4 with a mean tumour area of 14198,7 μm^2^ on D0 to 41159,6 μm^2^ on D4 (**** P-value < 0,0001) (*Figure 1D*). The tumour area was then normalized on D0 to assess the percentage of tumour growth (*Figure 1E*). Melanoma xenografts were characterized by an increase of tumour area of +254 % (** P-value = 0,0061) on D2 and +296 % on D4 (**** P-value = < 0,0001) (*Figure 1E*). This analysis demonstrated that melanoma xenografts exponentially and significantly increased their size in the first 2 days post transplantation to then grow at a slower rate.

**Fig.1:**
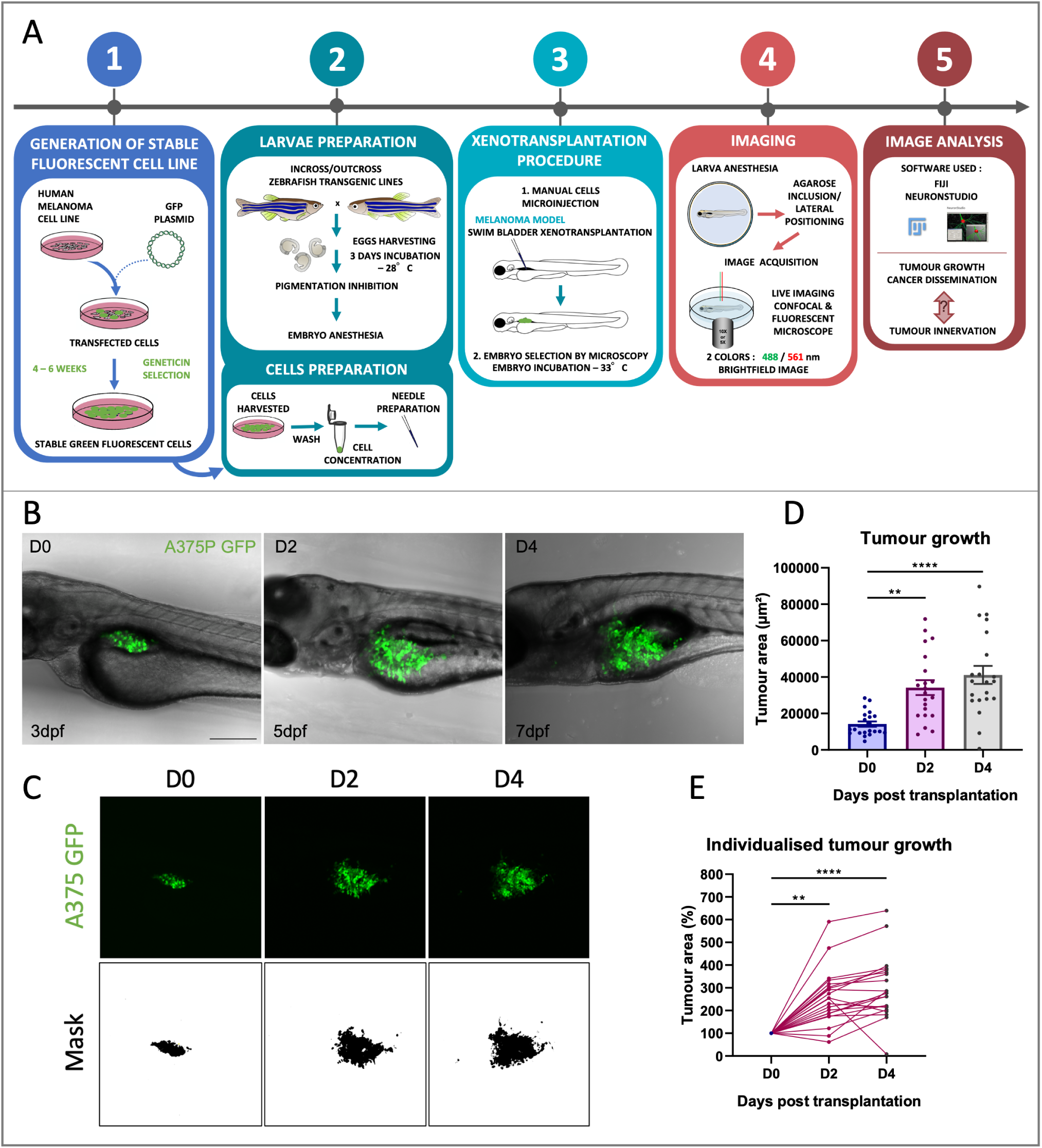
Evaluation of in vivo melanoma growth. **A.** Schematic anatomy of a 3 dpf zebrafish embryo. **B.** Transplanted larvae with A375P GFP+ (green) are shown in the images, acquired with a confocal microscope from D0 to D4. Images derive from Z-projection (z-stack = 170-200 μm). Scale bar = 250 μm. **C.** Representative xenografted melanoma tumours (green or black) are shown to explain how the analysis of tumour growth was made. Tumour growth was measured on the maximum projection of z-stack acquisitions. To determine tumour area, a threshold was applied in Fiji for each larva on D0, D2 and D4 to generate a mask. First line: maximum projection of the images on D0, D2 and D4 (z-stack = 170-200 μm). Second line: masks of the corresponding tumours, generated using Fiji software. **D-E**. Tumour growth analysis. The tumour area of individualized larvae was daily monitored from D0 to D4. The tumour area is expressed in μm^2^ (**D**). Individualised tumour areas were normalized to D0 to monitor the percentage of the tumour growth in every single larvae (**E**). Results in (**D-E**) are expressed as mean ±SEM, n = 21. Friedman test was used to evaluate the significance: * P-value<0,05, ** P-value <0,01, *** P-value <0,001, **** P-value < 0,0001.

To further characterize our xenograft melanoma model, we then focused on the morphology and the migratory capacity of A375P cells. The high invasive capacity of our melanoma model was assessed by counting the number of elongated and detached cancer cells from the primary tumour mass. Images showed the presence of morphological changes of melanoma cells (*Figure 2A, white arrowheads, C).* On D0 most of the A375P cells presented a round shape and few cancer cells were characterized by ellipsoidal morphology. On D2, but mostly on D4, several melanoma cells presented a very elongated morphology in comparison to D0 (*Figure 2A, white arrowheads*). Quantification analysis demonstrated a +492% significant increase of the number of elongated cells on D2 (** P-value = 0,0061) and +861% significant increase on D4 (**** P-value < 0,0001) in comparison to D0 (*Figure 2B*). These data showed a significant increase of elongated cells also between D2 and D4 (* P-value = 0,0129), demonstrating a continuous evolution towards a more infiltrative phenotype.

**Fig.2:**
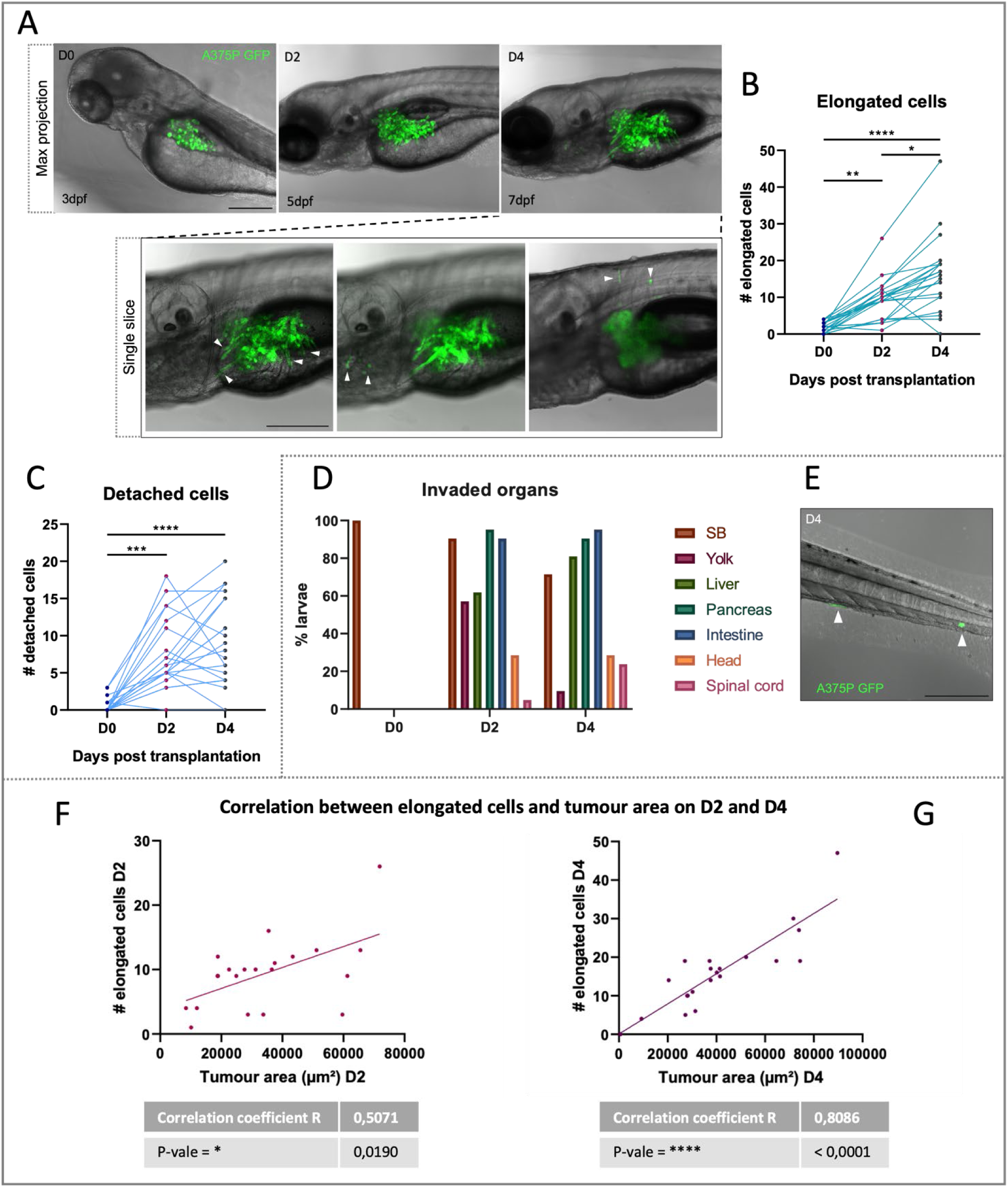
In vivo invasiveness and dissemination of A375P cells after xenografting in larval zebrafish. **A.** Representative images of transplanted larvae with invading and migrating A375 GFP+ cells (green) on D0, D2 and D4. First line: z-projection images (z-stack = 170-200 μm). Second line: three different single slices of the same image acquired on D4. Scale bar = 250 μm. White arrowheads point elongated and disseminated cells. **B.** The graph shows the quantification of elongated cells. The invasive cells were counted daily in every larva on D0, D2 and D4. **C.** The graph shows the quantification of detached cells from the primary cancer mass. The detached cells were counted daily in every larva on D0, D2 and D4. Friedman test was used to evaluate the significance in (**B**) and (**C**): * P-value<0,05, ** P-value <0,01, *** P-value < 0,001, **** P-value < 0,0001. n = 21. **D.** A375P cells invade the surrounding organs of the SB in larval zebrafish. The graph shows the percentage (normalized to the total number of larvae) of larvae with melanoma cells in the surrounding organs/regions near the SB on D0, D2 and D4. **E.** Representative image of the region of the tail of a larval zebrafish on D4. In green, A375P cells that migrate from the primary cancer mass to the CHT, in the tail. **F-G**: The graphs represent the correlation between the number of elongated cells and the tumour area (μm^2^) on D2 (E) and on D4 (F). Scale bar = 250 μm. Non-parametric Spearman correlation was performed to test the possible correlation: * P-value<0,0332, ** P-value <0,0021, *** P-value <0,0002, **** P-value < 0,0001. n = 21.

After transplantation, there was a significant number of disseminated cells on D2 (*** P-value = 0,0002) and D4 (**** P-value < 0,0001) in comparison to D0 (*Figure 2C*). Exploiting larval transparency, we then mapped the presence of cancer cells in the tissues and organs surrounding the SB, including in the yolk sac, liver, pancreas, intestine and in the region of the head and the spinal cord (*Figure 2D*) but also in distant locations, including the caudal region of the larvae (*Figure 2E*).

We then studied a possible correlation between the tumour area and the number of elongated cells on D2 and D4. On D2, the correlation coefficient R was higher than 0 (R = 0,5071, * P-value = 0,0190), indicating a direct relation between the two parameters (*Figure 2F*). This correlation was stronger on D4 (R = 0,8086, **** P-value < 0,0001) (*Figure 2G*). This significant correlation indicates that the number of elongated cells present in the larval body increases as the tumour growth, supporting the aggressiveness of our xenograft melanoma model.

To validate the presence of an aggressive melanoma, we performed gene expression analysis to check the immune response, hallmark of cancer ^32^ (*Sup.* Fig. 1). The expression of the two zebrafish pro-inflammatory cytokines *il1b* ^33^ and *il8* ^34^ was tested in the larval xenografts, also called melanoma transplants (MT), and in control larvae (CTL) injected with PBS in the SB. The analysis of the two zebrafish interleukins were significantly increased in MT in comparison to CTL, both on D0 (*Sup.* Fig. 1A *& 1C*) and D2 (*Sup.* Fig. 1B *& 1D*). We also compared the expression of human *IL8* in the MT on D2 and D4, which can be also released by cancer cells promoting pro-tumoral events, like angiogenesis ^34^. Interestingly, *IL8* expression was statistically reduced on D4 in comparison to D0 (*Sup.* Fig. 1E), indicating a possible reduction of the inflammatory status of the cancer niche. To compare these results with another zebrafish melanoma model, we checked the gene expression of *il1b* and *il8* in zebrafish melanoma biopsies, obtained from adult *kita:ras* zebrafish that spontaneously develop melanoma ^35^. *Kita:ras* biopsies had an higher *il1b* and *il8* expression in comparison to control skin biopsies, obtained from wild type adult zebrafish (*Sup.* Fig. 1F-G). Thus, zebrafish melanoma models seem to recapitulate the human condition of an inflammatory status. Together, our model recapitulates aspects of malignant human melanoma. Thus, we focused on identifying the presence of tumour innervation and its possible roles on cancer progression in this *in vivo* model.

### Tumour-induced axonogenesis in melanoma xenografts

To visualize the presence of tumour innervation in our melanoma xenograft model, A375P cells were transplanted in Tg(*nbt:DsRed)* transgenic zebrafish larvae ^36^ (*Figure 3A*) in which all neurons express the DsRed fluorescent protein. Confocal microscopy imaging revealed morphological changes of the motoneurons (*Figure 3A, white asterisks*) whose soma are localized in the spinal cord. Moreover, it was possible to discriminate the enteric neural cell bodies in the region of the intestine (*Figure 3A white arrowheads*). In addition, human melanoma cells (green) started invading the surrounding tissues by interacting with neurons (red) and migrating along their axons (*Figure 3B*, *white arrowheads*), in an event that resembles PNI ^9, 37^.

**Fig.3:**
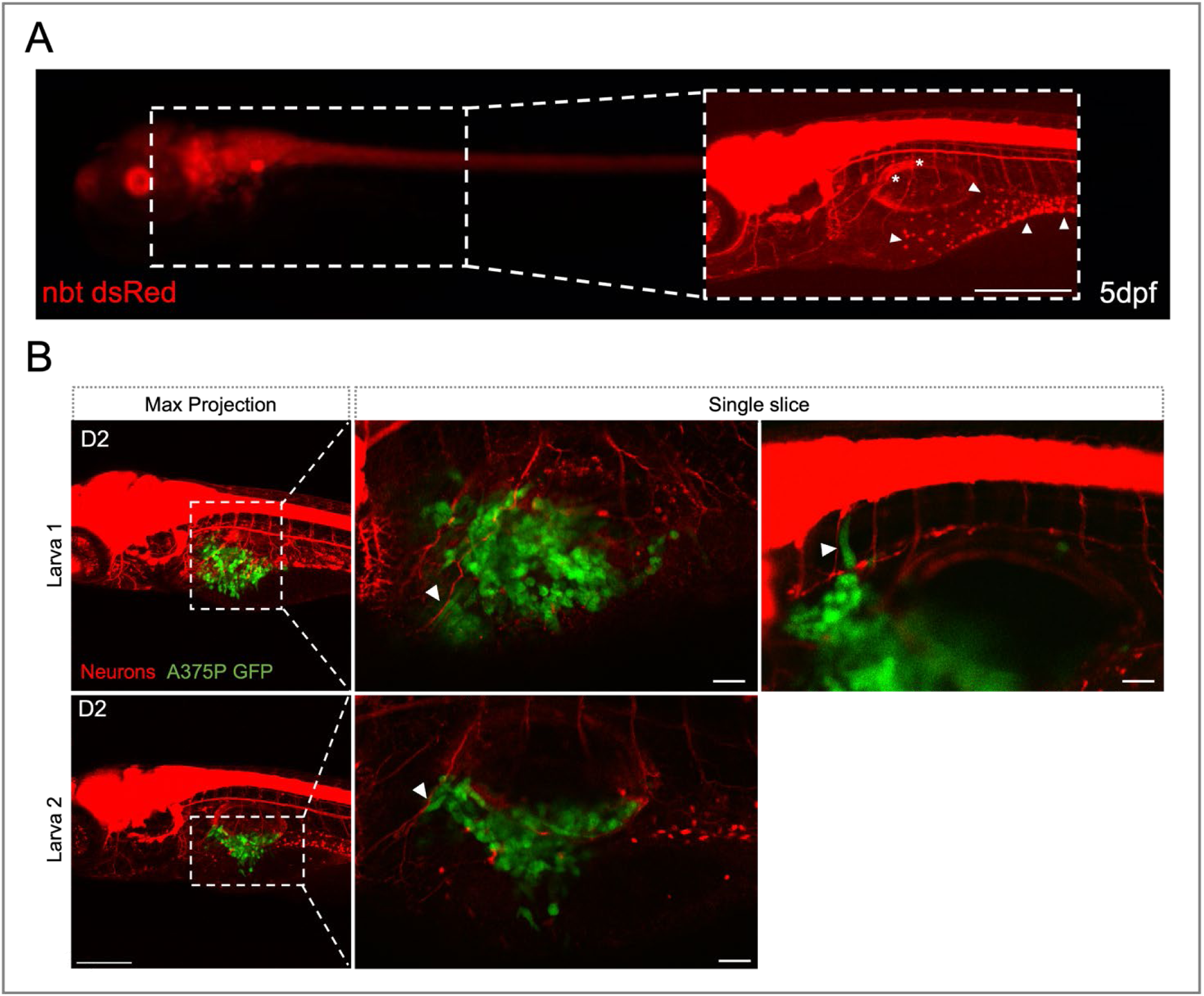
Interactions between PNS and transplanted melanoma cells in larval zebrafish. **A**. Representative images of a 5 dpf *tg(nbt:dsRed)* larva, acquired with a fluorescent microscope. Zoomed area: picture of the same larvae acquired with a confocal microscope. Scale bar = 250 μm. White arrowheads = enteric neural cell bodies. White asterisks = motoneurons. **B**. Representative images of xenografted larvae with A375P cells on D2, acquired with a confocal microscope. Transplanted A375P cells are in green, neurons are in red. A375 cells seem to escape from the primary cancer mass by following the axons as a route of dissemination (white arrowheads). First column: z-projection images (z-stack = 170-200 μm) of larva 1 and larva 2. Scale bar = 250 μm. Second and third columns: zoomed images of the region around the SB of larva 1 and larva 2 corresponding to different single slices. Scale bar = 50 μm.

To quantify potential changes of neural morphology, we focused on five axons located around the region of the nascent SB in the tumour niche (*Figure 4A*). First, we visualized interactions between melanoma masses and the five axons under study. The axons that were mostly involved in interactions with cancer cells were axons number 2 and 3 (*Figure 4*). We then measured axon length and monitored the number of dendritic branches. To evaluate the axons’ growth, we measured the length of the five axons on D0 and D4 in every single larva. Then, for every larva, the average of the five axons’ lengths was calculated (*Figure 4D*). On D0, there was no difference between the average axons’ length in CTL larvae, in comparison to those in the MT larvae. However, on D4 there was a significant increase in the average axons’ length in the MT larvae compared to the CTL larvae (*Figure 4D* * P-value = 0,0111). To evaluate which axons were the most affected by the presence of cancer cells, we performed the statistical analysis for each of the 5 axons in every larvae. On D4 the axons with a significant increase in length were axon number 2 (*Figure 4E*, * P-value = 0,0282) and 3 (*Figure 4F*, * P-value = 0,0301). This is correlated with the presence of a high number of cancer cells in the environment near these axons. The graphs related to axon number 1, 2, 3, 4 and 5 on D0 and axon number 1, 4 and 5 on D4 are reported in *Sup.* Fig. 2A and in *Sup.* Fig. 3A.

**Fig. 4:**
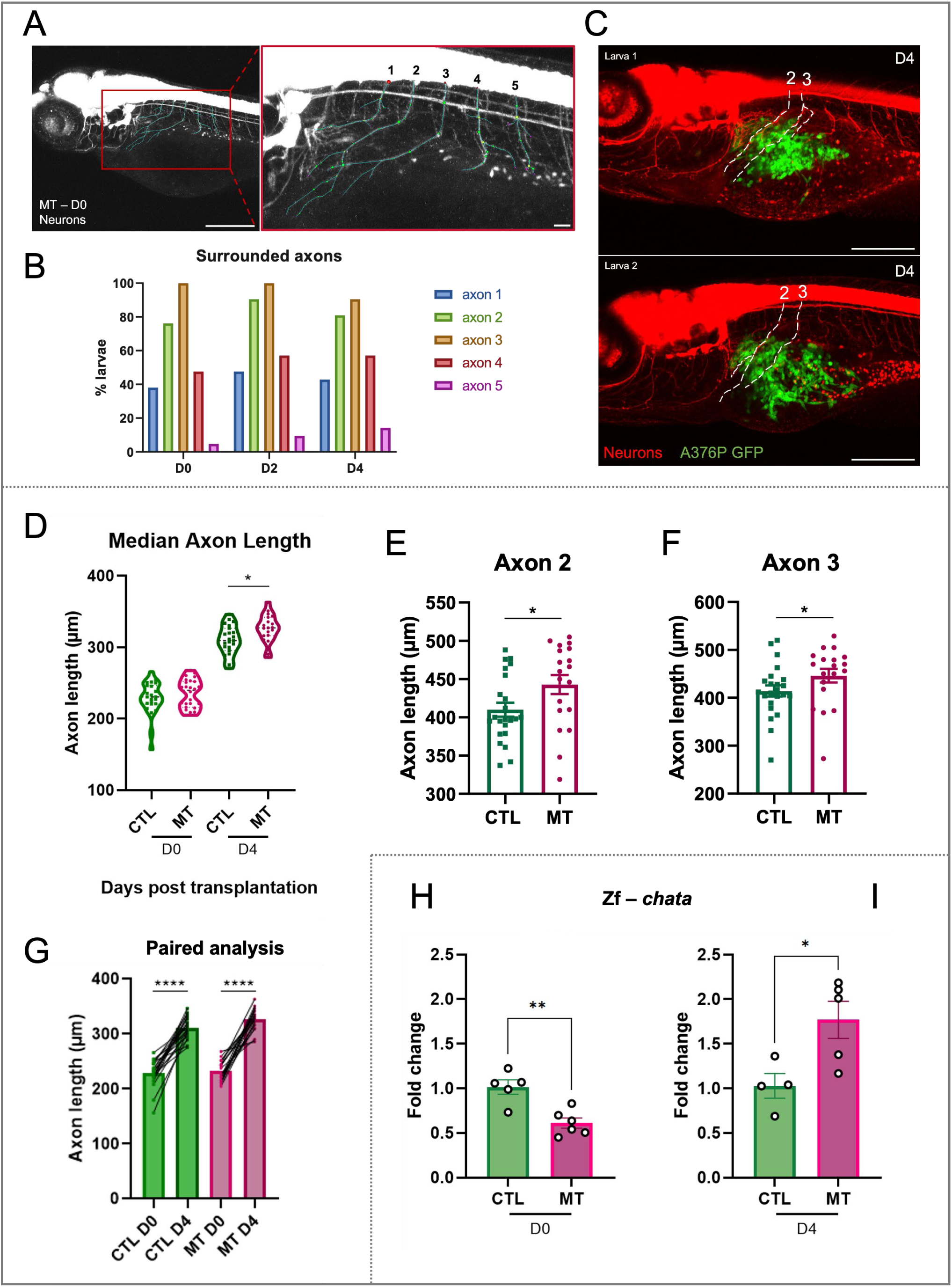
Axonogenesis in zebrafish xenograft melanoma model. **A**. Representative images of a 3 dpf larva from NeuronStudio software. Five axons around the region of the SB were selected and manually traced in 3D using NeuronStudio to measure the axon length and to count the dendritic branching points. Scale bar = 250 μm/ Scale bar = 50 μm. **B**. The graph shows the percentage of larvae with axons 1-5 surrounding A375P cells. **C**. Representative images of 7 dpf (D4) larvae with marked axon number 2 and 3. The images are taken using a confocal microscope. Scale bar = 250 μm. **D-G**. Axonogenesis was observed in the xenograft melanoma model. **D**. The graph shows the average length of the five axons in every single larva in every condition on D0 and D4. n=19-26. The measure of the axon length was performed using the NeuronStudio software. After manually tracing the axons and the dendritic branches, the software gave a summary with the length of every axon. **E-F**. The graphs show the axon length on D4 of the axon number 2 (**E**) and axon number 3 (**F**), the most affected axons by the presence of A375P cells. Differences among groups were analysed by two-tailed unpaired Student’s t test in (**D-F**): * P-value<0,05. n=19-23. **G**. The graph represents the paired analyses of the average of the five axons length of CTL and MT conditions on D0 and D4 as indicated. Differences among groups were analysed by two-tailed paired Student’s t test in (**G**): * P-value<0,05, ** P-value <0,01, *** P-value <0,001, **** P-value < 0,0001. n=19-22. The values in (**D-G**) are expressed in μm. **H-I**. Quantitative PCR analysis of the expression of the presynaptic cholinergic gene, choline acetyltransferase (*chata*), in the CTL and MT samples on D0 and D4 as indicated. Two-tailed unpaired Student’s t test was performed to evaluate the significance: * P-value<0,05, ** P-value <0,01. n=4-6. Results in (**E-F**) and (**H-I**) are expressed as means ± SEM. CTL = larvae injected with PBS MT (melanoma transplant) = larvae transplanted with A375P GFP + cells.

These axons belong to motoneurons, cholinergic neurons, that once completely mature and functional, produce and release the neurotransmitter acetylcholine. ChAT, choline acetyltransferase, catalyses the rate-limiting step in the acetylcholine biosynthesis. Due to zebrafish genome duplication, this enzyme is encoded by two genes, *chata* and *chatb*, that are differentially expressed in different types of neurons. However, generally, *chata* is the only one expressed in early larval stage ^38^ as also demonstrated by our gene expression analysis. Indeed, the expression of both genes were examined by real-time PCR, but the analysis demonstrated detectable levels only for the *chata* gene.

On D0, about 6h post transplantation, there is significant decrease of *chata* expression level in MT samples in comparison to CTL larvae (*Figure 4H*, ** P-value = 0,0023). This could be linked to the possible stress or neuron-injury associated to the injection of human cancer cells in the larval SB. On the contrary, on D4, the presence of human cancer cells increased *chata* expression in MT samples in comparison to the CTL larvae (*Figure 4I*, * P-value = 0,0265). This can be correlated to a higher activity of the motoneurons and other cholinergic neurons in the larval zebrafish, such as the subpopulation of enteric neurons ^39^.

Together, these results support the presence of axonogenesis in the TME of our larval zebrafish xenograft melanoma model.

### Tumour-induced dendritogenesis in melanoma xenografts

We also examined the number of dendritic branching points as a readout of dendritogenesis, in the context of tumour innervation. We mapped the dendritic arborization of the five selected axons, using the NeuronStudio software and found that already from D0, 6h post transplantation, there was a significant increase (** P-value = 0,0033) of the average number of dendritic branching points in MT larvae in comparison to the CTL (*Figure 5A, 5F*). The most affected axons were axons number 2, 3 and 5 (*Sup.* Fig. 2B). This increase became bigger on D4 (*Figure 5A*, **** P-value < 0,0001), and in this case all the 5 axons of the transplanted larvae had an increased number of dendritic branching points in comparison to the CTL larvae (*Figure 5B, Fig. Sup. 3B*). This indicates that A375P cells provide trophic support for neurite’s growth inside the TME. To complete the analysis of the dendritic morphology, we performed Sholl analysis (*Figure 5C*). The data acquired from this analysis allowed the construction of the Sholl profile (*Figure 5D*, *Sup.* Fig. 2C *& 3C*). On D0, the Sholl profiles of the 5 axons did not show significant differences between the CTL and the MT larvae, even though there was a tendency to increase in the MT samples (*Sup.* Fig. 2C). On D4, the MT Sholl profiles exhibited a higher increase in comparison to the CLT ones (*Sup.* Fig. 3C); the MT profile with a significant difference from the CTL one is the one for axon 2 (*Figure 5D*). Indeed, all the points related to the MT curve are positioned higher in comparison to the CTL curve. This means that the most affected axon by the presence of human A375P cells is the number 2, both in the increase of the axon length (*Figure 4E*), the increase of the number of dendritic branching points (*Figure 5B*) and the Sholl profile (*Figure 5D*).

**Fig.5:**
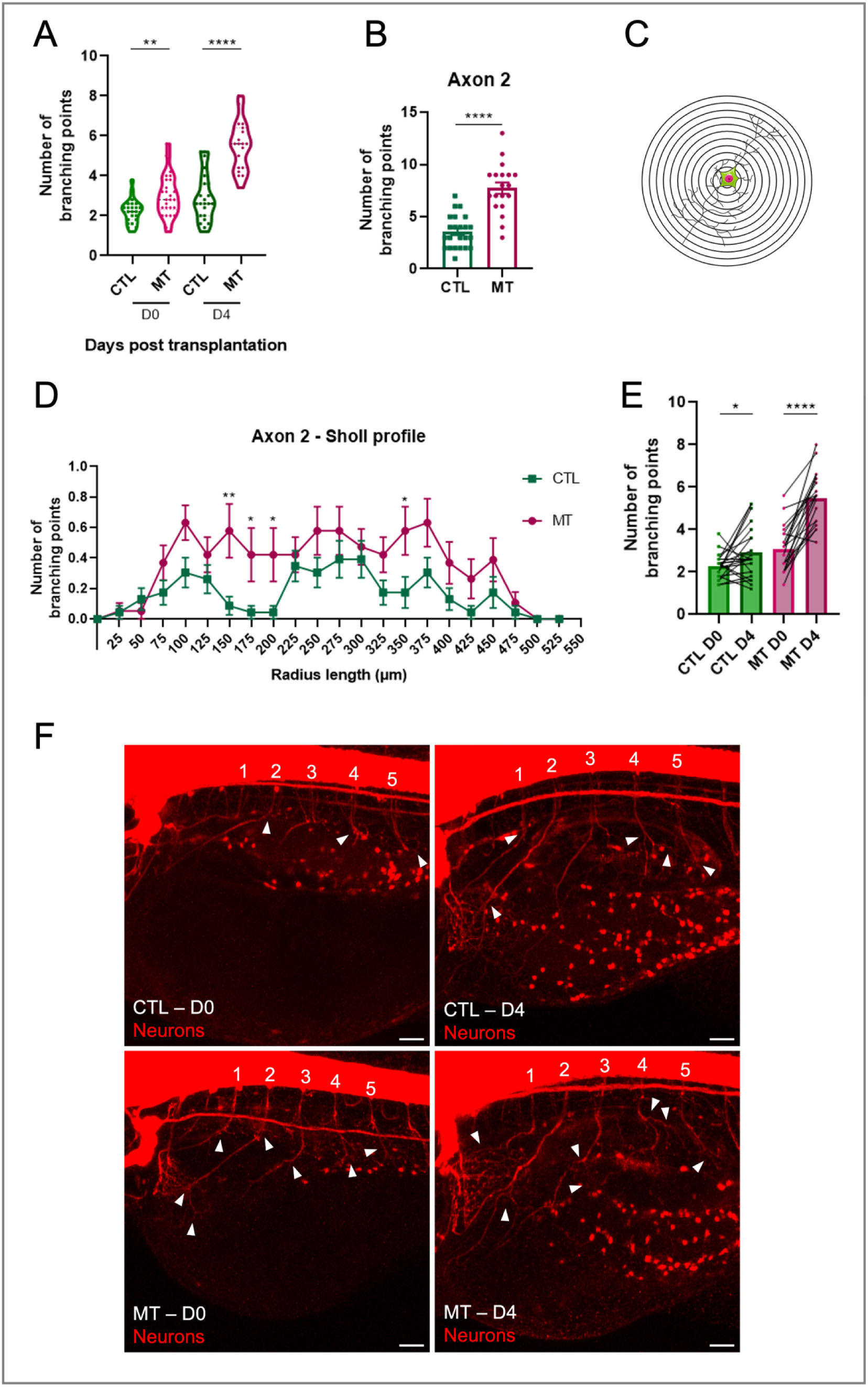
Dendritogenesis in zebrafish xenograft melanoma model. **A.** The graph represents the average of the branching points’ number of the five axons in every larva in CTL and MT conditions on D0 and D4. n=19-26. **B.** The graph represents the number of branching points of the axon number 2 in every larva in CTL and MT conditions on D4. n=19-23. Results are expressed as means ± SEM. Differences among groups were analysed by two-tailed unpaired Student’s t test in (**A-B**): * P-value<0,05, ** P-value <0,01, *** P-value <0,001, **** P-value < 0,0001. **C.** Schematic view that allows the description of the Sholl analysis, a quantitative method to study the dendritic anatomy. The picture shows concentric rings centred on the soma centre of the neurons. From the centre, the Sholl analysis starts counting the number of the intersections of the dendrites with the different rings, reporting the distance from the soma centre. The number of intersections is the number of the dendritic branches. The data acquired from the analysis allow the construction of the Sholl profile. **D.** The Sholl profile of axon number 2 on D4. The sholl profile is a graph that plots the number of the branching points against the radial distance from the soma centre. The radial step between every ring is 25 μm. The axons analysed in this work are from motoneurons, whose somas are placed in the spinal cord. Thus, the soma centre was generally placed in the ventral part of the spinal cord for every neuron/axon. Differences among groups were analysed by Mann-Whitney test: * P-value<0,05, ** P-value <0,01. n = 22. Results in (**B**) and (**D**) are expressed as means ± SEM. **E.** The graph represents the paired analyses of average of the five axons’ branching points of CTL and MT conditions on D0 and D4 as indicated. Differences among groups were analysed by two-tailed paired Student’s t test in (**D-F**): * P-value<0,05, ** P-value <0,01, *** P-value <0,001, **** P-value < 0,0001. **F.** Representative images of nbt CTL and MT larvae on D0 and D4. Neurons are in red. White arrowheads point the branching points. Images are taken using a confocal microscope. Scale bar = 25 μm. CTL = larvae injected with PBS MT (melanoma transplant) = larvae transplanted with A375P GFP + cells.

As PNS developments is still ongoing in the larval stage, we performed an additional analysis from our data in which the evolution of the average of the axons’ length and dendritic branching points in every single larva on D0 and D4 was considered. Paired analysis of CTL and MT showed an increase of branching points on D4 in the transplanted embryos in comparison to the CTL (*Figure 5E*; CTL: * P-value = 0,0335 VS MT: **** P-value < 0,0001), despite the presence of the developmental force. This difference is not so visible focusing on the measure of the axon length (*Figure 4G*; CTL: **** VS MT: ****, P-value < 0,0001). This indicates that the presence of cancer cells impacts more on the formation of new branching points, than on the promotion of axon length where the PNS developmental force is much more relevant.

### Tumour-induced neurogenesis in melanoma xenografts

Due to the high invasion of the intestinal tract by the A375P cells, we monitored the enteric neurons to visualize a possible effect of melanoma cells on this specific cell type. The zebrafish neural progenitors of the enteric neurons populate the intestinal epithelium between 32 hours post fertilization (hpf) and 66 hpf ^40^ and the intestinal tract is clearly recognizable from late 3 dpf. Thus, thanks to the *nbt* transgenic line, this neural population is detectable inside the intestinal area (*Figure 3A & 6A*, *white arrowheads*). Quantitative *in vivo* image analysis demonstrated a significant increase of the number of the enteric neural cell bodies in the MT larvae in comparison to the CTL larvae on D4 (*Figure 6A & 6B,* * P-value = 0,0293) but not on D2. This analysis shows that tumour xenografts induce neurogenesis in the region of the intestinal tract. To validate this result, we quantified the expression of two genes, *elavl3* and *sox10*, important in the development of the PNS. *Elavl3* gene encodes for a protein that is a marker of early neurogenesis ^41^ ^42^, and consistent with an induced neurogenesis, its expression was significantly increased in xenografted larvae in comparison to the CTL (*Figure 6C-D*, D0 -* P-value = 0,0378, D4 -* P-value = 0,0161). Moreover, comparing its expression from D0 and D4 in MT samples, we observed an increase of e*lavl3* expression over the time.

**Fig.6:**
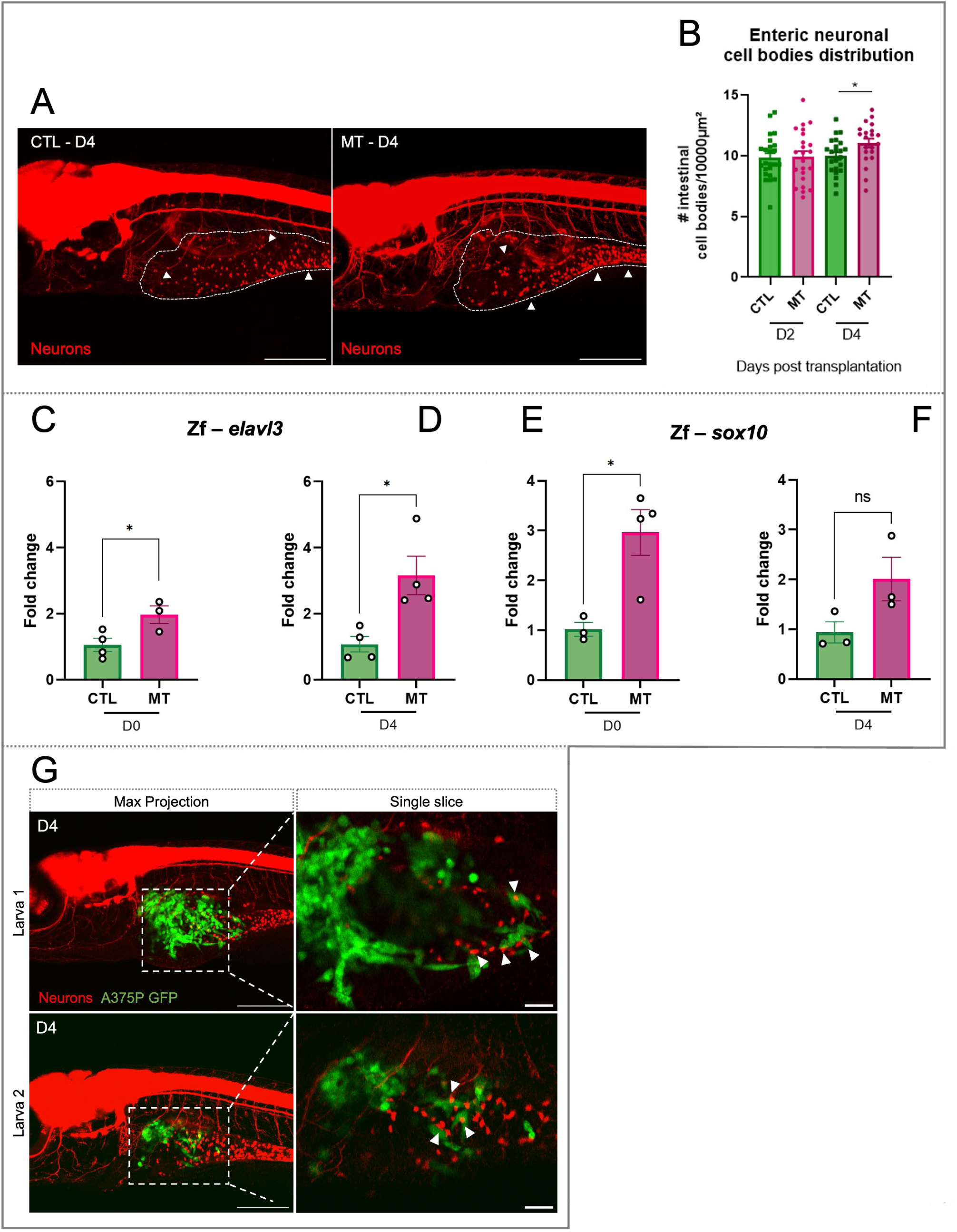
Neurogenesis in zebrafish xenograft melanoma model. **A.** Representative images of *nbt* CTL and MT larvae on D4. Neurons are in red. Surrounded areas delimit the intestinal zone, white harrowheads point the enteric neural cell bodies. Images are taken using a confocal microscope. Scale bar = 250 μm. **B.** The graph shows the number of enteric neural cell bodies per 10^4^ μm in CTL and MT conditions on D2 and D3. Differences among groups were analysed by two-tailed unpaired Student’s t test in: * P-value<0,05. n=21-24. **C-F.** Quantitative PCR analysis of the expression of the early neurogenesis marker *elavl3* (**C-D**) and the NC marker *sox10* (**E-F**) in CTL and MT samples on D0 and D4 as indicated. Two-tailed unpaired Student’s t test was performed to evaluate significance: * P-value<0,05. n=3-4. Results in (**B-F**) are expressed as means ± SEM. **G**: Representative images of xenografted larvae with A375P cells on D4. Images were acquired with a confocal microscope. Transplanted A375P cells are in green, neurons are in red. First column: z-projection images (z-stack = 170-200 μm) of larva 1 and larva 2. Scale bar = 250 μm. Second column: single slice of the zoomed area of the corresponding projection images. White arrowheads point the contact between enteric neural cell bodies and cancer cells. Scale bar = 50 μm. CTL = larvae injected with PBS MT (melanoma transplant) = larvae transplanted with A375P GFP + cells.

*Sox10* is a neural crest marker and transcription factor important for the development of enteric neural progenitors ^43^, Schwann cells ^44^ and melanocytes ^45^. Its expression was higher in the xenografted larvae in comparison to the CTL, even though the result is statistically significant just on D0 (*Figure 6E-F*, * P-value = 0,0170). High *sox10* expression can be associated to the enteric neurogenesis process and at the same time to an increase of the Schwann cells in the MT larvae. The expression of *sox10* was also checked in the *kita:Ras* melanoma biopsies. In this case, the increased level of *sox10* transcript was associated to the presence of the zebrafish melanoma mass, which is completely absent in the wild type (CTL) biopsies (*Sup.* Fig. 1H).

Interestingly, in high resolution images, we also observed contacts between A375P melanoma cells and enteric neural cell bodies (*Figure 6G, white arrowheads*). The direct contact could mediate a reciprocal communication between the two different cell types, supporting, on one hand, cancer invasiveness, and, on the other hand, tumour innervation through an increase of neurogenesis. Altogether these results demonstrate the engagement of the PNS in melanoma xenografts.

### Catecholamines in in vivo xenograft model support melanoma progression

We then characterized the role of innervation on cancer progression. Many studies have reported that the autonomous PNS is involved in cancer progression ^14,27,46^. Focusing on malignant melanoma, *in vitro* and clinical data suggest that adrenergic signalling are relevant in melanoma proliferation and invasion ^24,25^ ^26,27,28^. To investigate this *in vivo*, we injected A375P cells with 10 µM noradrenaline (NA) or 1 µM adrenaline (AD), mimicking the presence of catecholamines in the TME, to study their impact in our xenograft melanoma model. The concentrations of noradrenaline and adrenaline were chosen according to literature ^47,48,49^. Initial concentration of 10 µM was used both for NA and AD, however *in vitro* AD treatment of A375P cells inhibited cell growth (*data not shown*), thus its concentration was reduced by 10 times. After transplantation, we examined the expression of the human *ADRB2* gene, that codifies for the β2-receptor. In the xenograft model, the expression was maintained from D0 to D4 at the same levels (*Sup.* Fig. 4A).

Transplanted larvae were monitored using live image acquisition on D0 and D3 to assess cancer growth, cell migration and invasive capacity. Catecholamines did not induce changes on tumour growth (*Sup.* Fig. 4B) but A375P cells had a significant higher migration capacity in the presence of NA (*Figure 7A*, * P-value = 0,0370, *Sup.* Fig. 4D). Moreover, in our *in vivo* model, catecholamines induced cells delamination associated with higher invasion of the tail (*Figure 7B &* *Sup.* Fig. 4D). As previously mentioned, our model is characterized by the presence of A375P cells migrating from the primary mass to distant sites, such as the region of the CHT (*Figure 2E*), thus we quantified tail’s invasion (*Figure 7C*) on D3. We observed a significantly increase (** P-value = 0,0097) of tail’s invasion by A375P cells in NA treated larvae in comparison to CTL ones (*Figure 7B*). AD also increased the number of larvae with invaded tail in comparison to CTL, supporting a tendency to increased cancer aggressiveness mediated by catecholamines. Finally, by looking at the morphology of the A375P cells, we found that catecholamines’ treatments induced an increased number of elongated cells on D3, however the results are not statistically significant (*Fig. Sup. 4C*).

**Fig.7:**
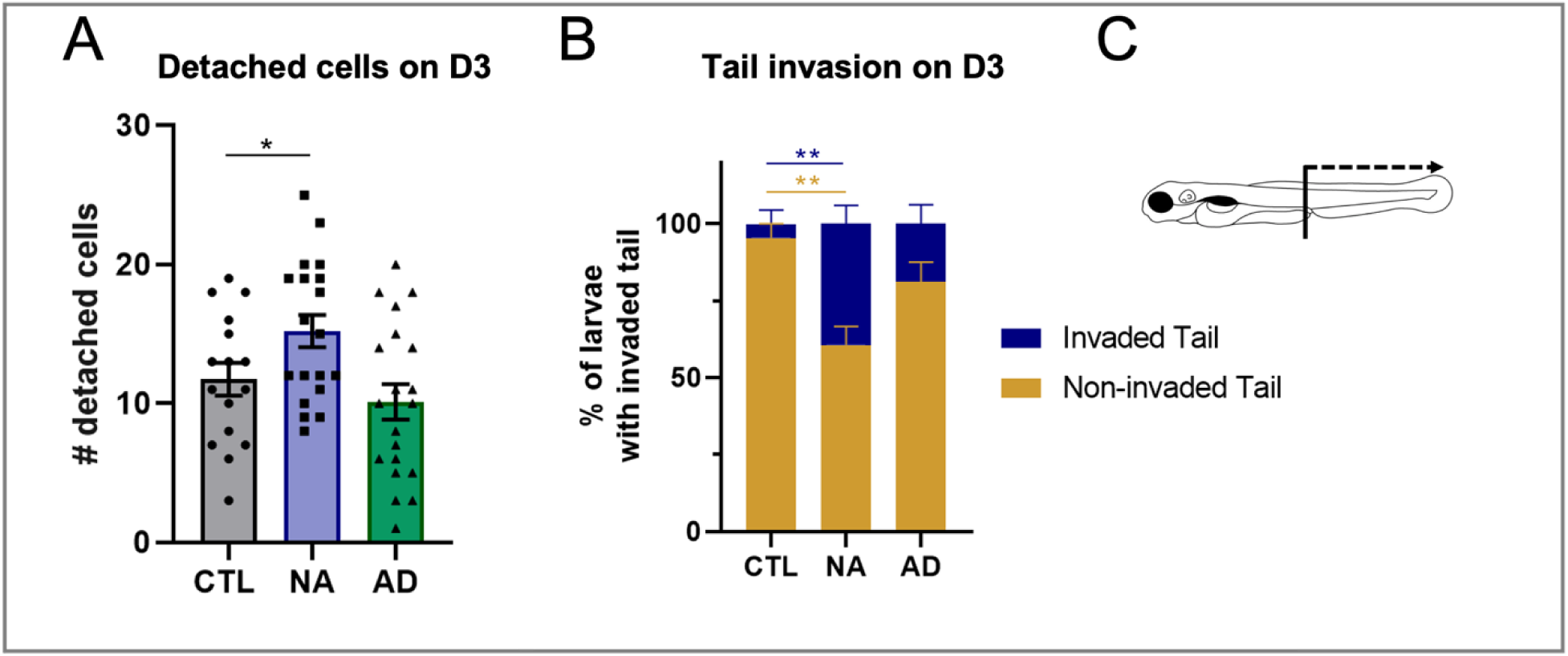
Effect of catecholamines in xenograft melanoma model. *In vivo* tumour invasiveness analysis upon co-injection with catecholamines. **A.** The graph shows the quantification of detached cells from the primary cancer mass on D3. Detached cells were counted daily in larvae for every condition on D0 and D3. Larval xenografts with already disseminated cells on D0 were excluded from the analysis, to avoid considering larvae with accidentally transplanted cells in the blood circulation. Differences among treated conditions and control condition were analysed by ordinary one-way ANOVA: * P-value<0,05. n=16-20. **B.** Graph represents the percentage of larvae with tail invaded/non-invaded by A375P cells in the three indicated conditions. Differences among treated conditions and control condition were analysed by two-way ANOVA: * P-value<0,0332, ** P-value <0,0021. n=19-20. **C.** Schematic representation of larva where the region of the tail is indicated. Cancer cells in the tail = detached cells localized from urogenital opening until the posterior part of the CHT. NA 10 μM = larvae transplanted with A375P cells and co-injected with 10 μM of noradrenaline. AD 1 μM = larvae transplanted with A375P cells and co-injected with 1 μM of adrenaline.

Together, these results demonstrate that catecholamines, especially noradrenaline, promote melanoma progression through increased cancer cell migration and invasion of distant sites.

## Discussion

In the present study, we have used in vivo imaging to characterize the impact of melanoma cells o nearby population of peripheral neurons. In addition, we show the contribution of catecholamines to melanoma progression towards a more aggressive disease. Performing *in vivo* analyses, we demonstrated the role of catecholamines in the progression of melanoma toward a more aggressive disease. This was possible thanks to the development of an *in vivo* melanoma model based on a novel larval zebrafish xenograft tool.

Transplantation of A375P human melanoma cells was associated with the development of a highly proliferative and invasive tumour mass, supported by the presence of an inflammatory TME. Thus, the zebrafish SB appeared as an optimal cancer niche for melanoma progression. Transplanted human melanoma cells could infiltrate into the surrounding organs, and also invade distant tissues. Invasion was compatible with the activation of the EMT-like process, also called phenotype switching in melanoma biology ^50^. Indeed, changes in melanoma cell morphology were clearly observed in our melanoma model. In detail, before and during the detachment of A375P cells from the tumour mass, the cells became elongated. However, most of them, when they were completely detached, were characterized by a rounder morphology. This phenomenon although needs to be further characterized demonstrates that the zebrafish TME is relevant to induce changes on human cancer cells, promoting an aggressive type of melanoma.

Transplanted A375P cells in larval zebrafish migrated specifically in the region of the tail where the CHT is located. Indeed, the CHT is a special niche, where a large panel of different secreted factors, such as the SDF1 ^51^, can attract cancer cells and sustain their survival and proliferation ^52,53^. The development of a very aggressive type of *in vivo* melanoma was fundamental for the following studies focused on the active function of innervation and catecholamines in melanoma progression.

In the last years, tumour innervation has become a novel hallmark of cancer initiation and progression, however until now only *in vitro* and mouse models were used to study it ^9, 19,20–21^. Here we demonstrated the high potential of the zebrafish model for *in vivo* studies of tumour innervation. Thanks to the availability of the *nbt* zebrafish transgenic line ^36^, we were able to visualize and follow in real time changes of the PNS in the TME of melanoma xenografts *in vivo*.

Here, we report the response of PNS to the presence of melanoma in terms of neurogenesis, axonogenesis, and dendritogenesis. While in physiological conditions, neurogenesis mostly occurs during embryonic development, early in the postnatal brain and in the adult enteric NS ^54,55^, it is reactivated in some malignancies, including prostate cancer, gastric and colorectal carcinoma ^56,57,58,59^. However, the precise molecular and cellular mechanisms governing this process have not been yet clarified. In our larval xenograft model, we confirm the presence of an active neurogenesis at the level of the enteric NS after detecting an increased number of enteric cell bodies in the intestinal tract, also supported by an increased expression of *elavl3*, an early marker of neurogenesis.

Moreover, similar to internal organs and tissues, several solid cancers are innervated by nerve fibres of the PNS, especially those arising from the autonomic NS, contributing to the formation of the TME ^14,16,19–21^. The activation of axonogenesis and dendritogenesis processes resemble the ones happening in the peripheral nerves after injury ^11^. In our model, we detected modifications in the morphology of the motoneurons localized in the region surrounding the melanoma mass, including increase in axonal length and in the number of dendritic branching points, generating a more complex dendritic arborization in the melanoma TME. High level of *chata* gene expression confirmed an increased cholinergic activity of the motoneurons but also of the cholinergic population of the enteric neurons.

In the adult PNS, changes in the neural network can be induced by Schwann cells, as they secrete several growth factors that can sustain axonal regeneration ^46^. Interestingly, cancer cells can secrete the so called cancer-related neurotrophic factors, that can similarly stimulate the growth of the axons and neurites in the TME ^37^. Once released, they create a molecular gradient that, working in an autocrine or paracrine manner, can promote both cancer cells proliferation and tumour innervation ^37,46^. Thus, tumour innervation is directly related to a mutual communication between cancer cells and the neurons in the tumour niche. In our model, we have shown an increase of the dendritic arborization from 6h post transplantation (*Graphical Abstract [1]*), compatible with A375P melanoma cells inducing dendritogenesis. We have also observed direct interaction between A375P melanoma cells and enteric neural cell bodies and observed that axonal ramifications were used as a route of cancer dissemination (*Graphical Abstract [2a][3]*). Indeed, elongated A375P cells were found following axons, in a way that resembles the PNI described in highly innervated cancers, similar to prostate and gastric carcinoma ^9,37^. PNI is reminiscent of a well described process, generally referred to as vessel co-option, that involves the vasculature system as a privileged route for cancer cell migration ^60^. Peripheral nerve fibres and vessels grow simultaneously and following the same pathways during development ^61,62^. Thus, although cancer research has devoted many efforts to the study of vessel co-option, further experiments, including denervation strategies, will help to further understand the impact of neurons as an additional route of metastatic dissemination. Moreover, during PNI, in addition to neurons being attracted by cancer cells, the axons can also exert a trophic function, as nerve endings can secrete factors supporting a positive microenvironment for the survival and proliferation of cancer cells ^63,64^. In this context, further studies will be necessary to characterize the molecular signalling controlling melanoma innervation and the identification of soluble factors that can help in the design of innovative anti-neurogenic therapies that specifically target the neural signalling inside the TME.

*In vitro* and clinical data suggest a role of adrenaline and noradrenaline, neurotransmitters from the sympathetic NS, in the regulation of melanoma progression ^24,27,65^. Thanks to the expression of the adrenergic receptors, human melanoma cells can respond to the catecholamines inside the TME. However, in zebrafish, the development of the sympathetic system happens between 2 dpf and 8 dpf ^42^, thus in our xenograft model it was difficult to study the direct contribution of the sympathetic system to melanoma progression. So, we co-injected A375P melanoma cells with either adrenaline or noradrenaline, to mimic the presence of catecholamines in the human TME. Our *in vivo* study has demonstrated that catecholamines, especially noradrenaline, promote melanoma progression in terms of increased cancer migration and invasion (*Graphical Abstract [3]*). This models an early-medium stage of melanoma progression, where A375P cells, escaping from the primary mass, probably undergo an EMT-like process. Our data suggest that catecholamines would enhance this behaviour.

Further experiments, that would deepen our knowledge of the role of catecholamines in tumour progression, include the assessment of the inhibitory activity of β-blockers on melanoma proliferation and metastatic colonization, for which the xenograft model could be appropriate as it offers the possibility to easily perform high-content drug screening. Interestingly, we have shown that melanoma cells have an impact on the neural microenvironment and that the use of a novel zebrafish preclinical model, which allows to visualize and monitor the dynamic interactions of the tumour cells with the neural microenvironment, will help to characterize the mechanisms by which the PNS regulates melanoma biology.

## Supplementary materials

### Tables

**Table 1:**
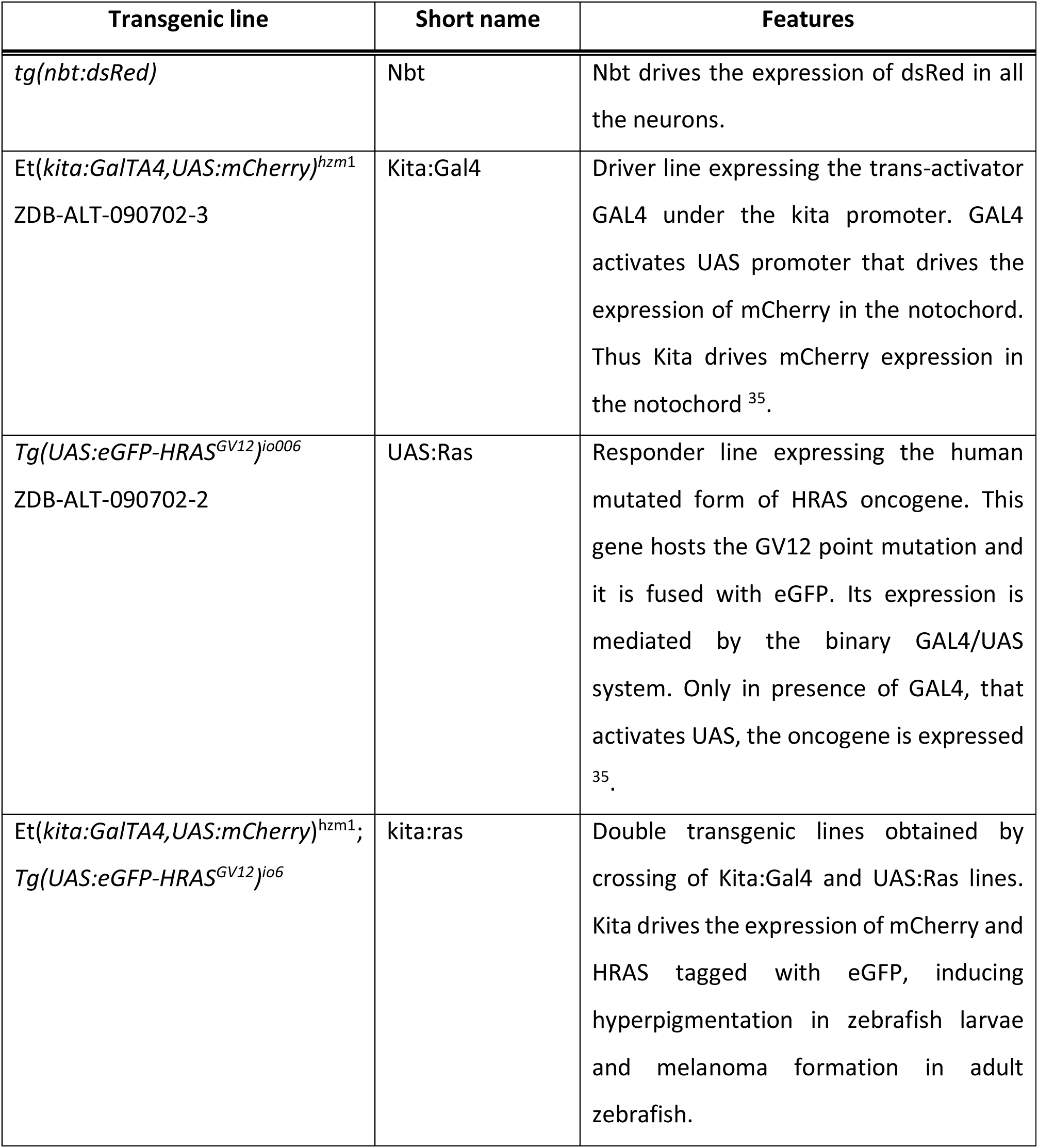
List of used transgenic lines and their features.

**Table 2:**
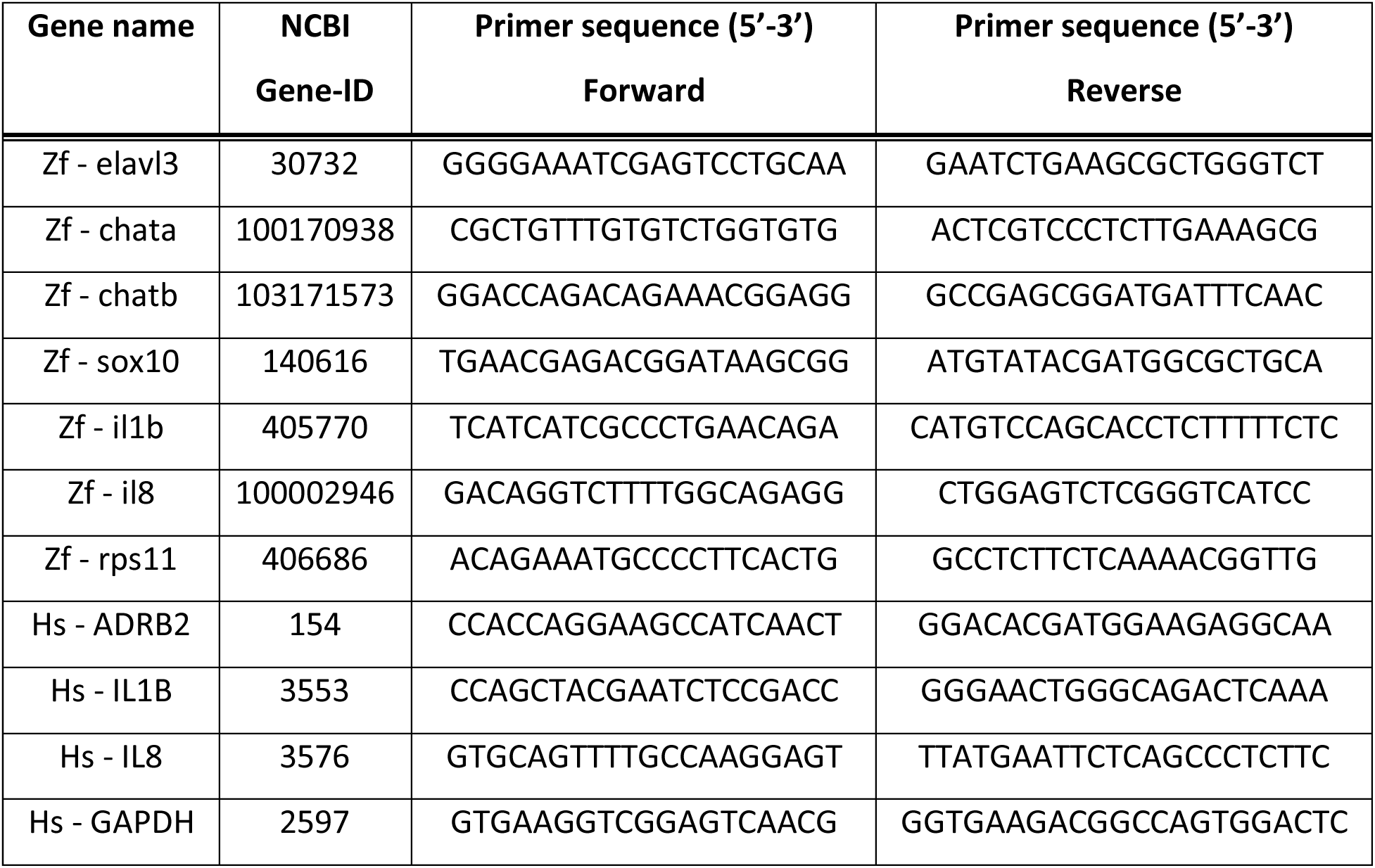
List of the qPCR primer sequences.

## Supplementary Figures

**Sup. Fig. 1:**
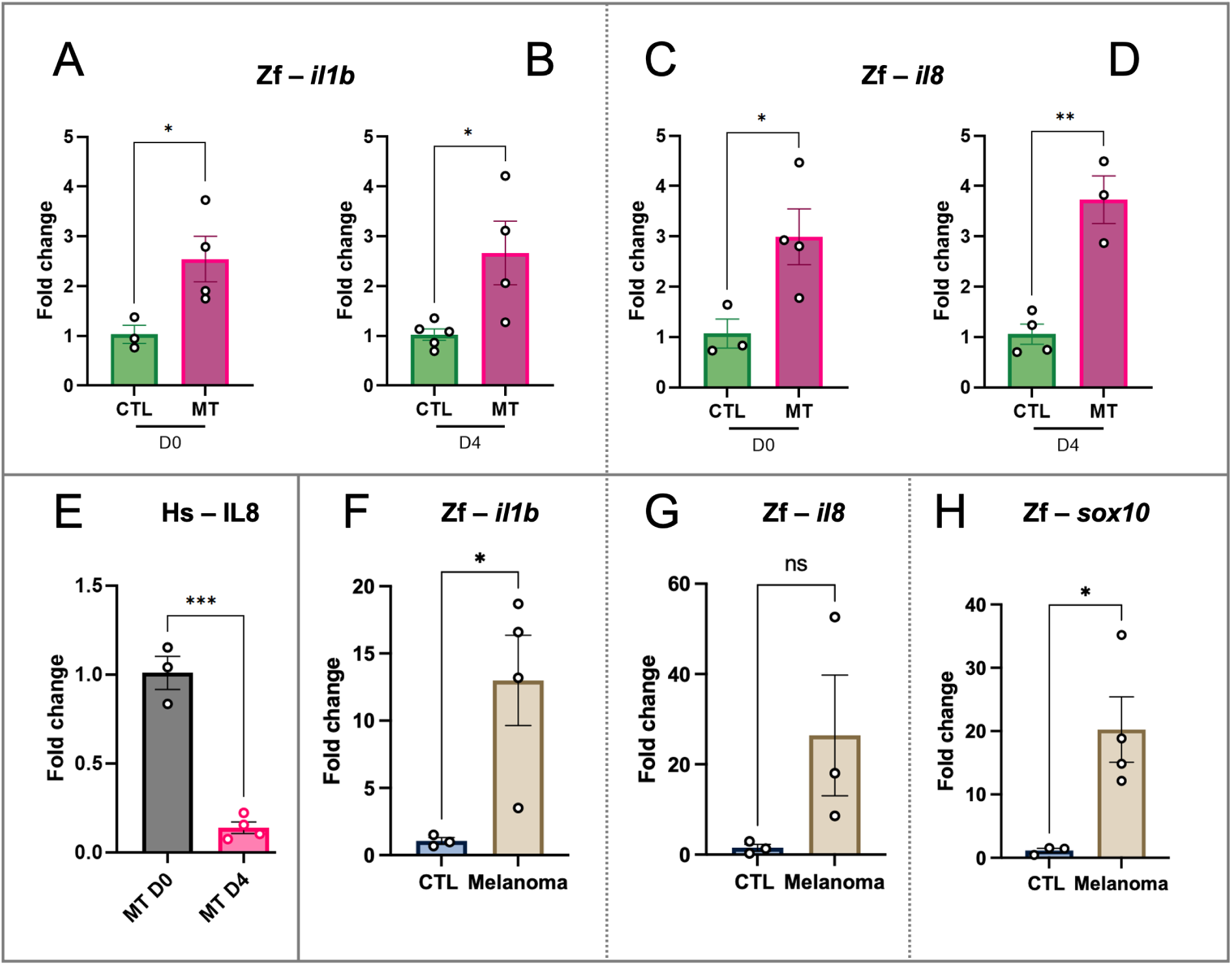
Gene expression analysis of pro-inflammatory markers and the NC marker sox10. **A-E.** Quantitative PCR analysis of the expression of zebrafish and human pro-inflammatory markers *il1b* (**A-B**) and *il8* (**C-D, E**) in CTL and MT samples on D0 and D4 as indicated. CTL = larvae injected with PBS MT (melanoma transplant) = larvae transplanted with A375P GFP+ cells. **F-H**. Quantitative PCR analysis of the expression of zebrafish pro-inflammatory markers *il1b* (**F**) and *il8* (**G**) and the NC marker *sox10* (**H**) in CTL and melanoma biopsies from respectively adult wild type and *kita:Ras* zebrafish. CTL = body biopsies of wild type adult zebrafish Melanoma = melanoma biopsies from adult *kita:ras* adult zebrafish Results (**A-H**) are expressed as means ± SEM. Two-tailed unpaired Student’s t test was performed to evaluate the significance: * P-value <0,05, ** P-value <0,01, *** P-value <0,001. n=3-4.

**Sup.Fig.2:**
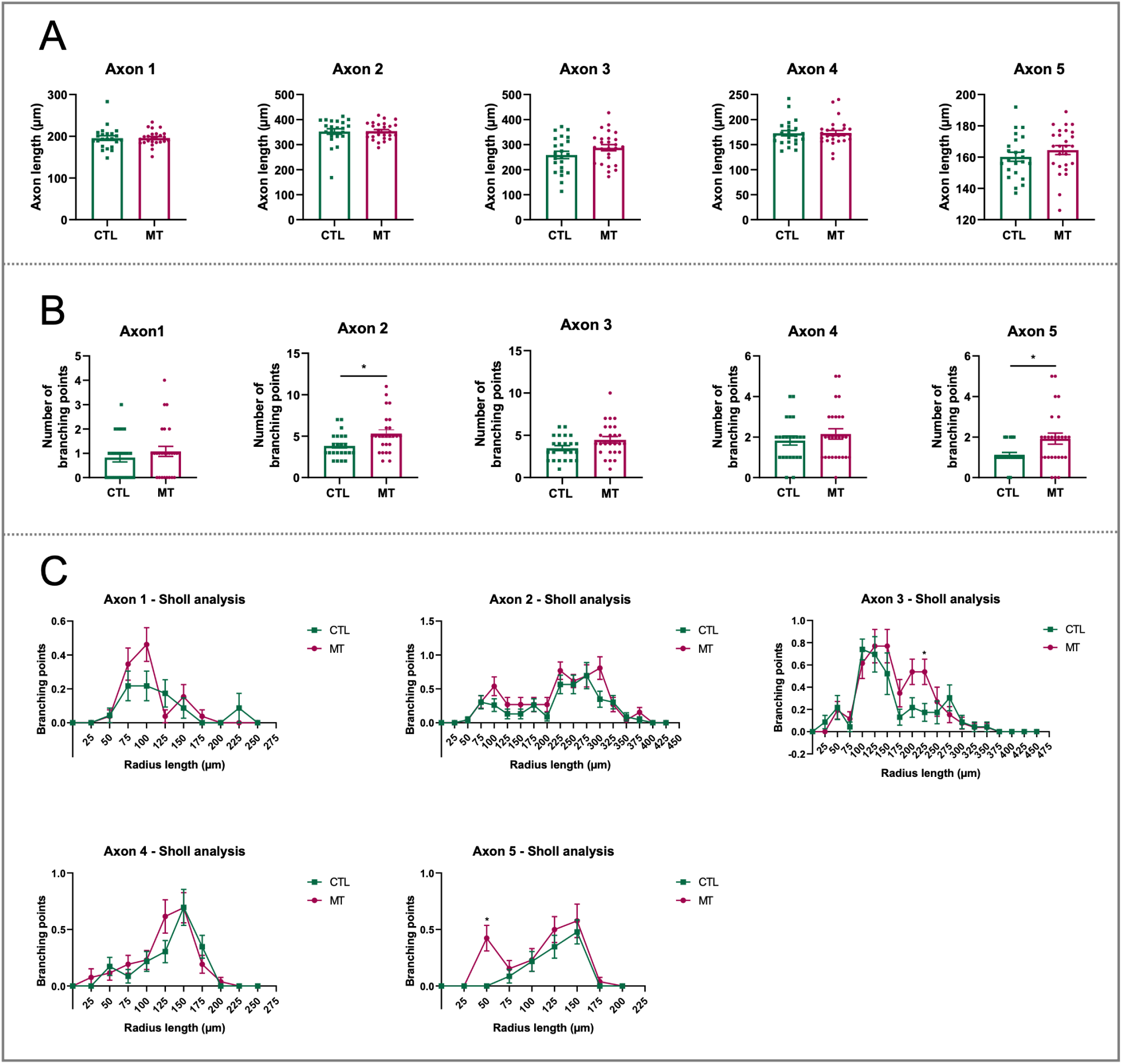
Complete tumour innervation analysis on D0. **A.** The graphs show the axon length on D0 of the axon number 1, 2, 3, 4 and 5 of CTL and MT larvae as indicated. The axon length is expressed in μm. **B.** The graphs represent the number of branching points on D0 of the axon number 1, 2, 3, 4, and 5 of CTL and MT larvae as indicated. Differences among groups (**A-B**) were analysed by two-tailed unpaired Student’s t test: * P-value<0,05, ** P-value <0,01. **C**: Sholl profiles of axon number 1, 2, 3, 4, and 5 on D0 of CTL and MT larvae. Differences among groups were analysed by Mann-Whitney test. Results are expressed as means ± SEM. n=23-26. CTL = larvae injected with PBS MT (melanoma transplant) = larvae transplanted with A375P GFP + cells.

**Sup. Fig. 3:**
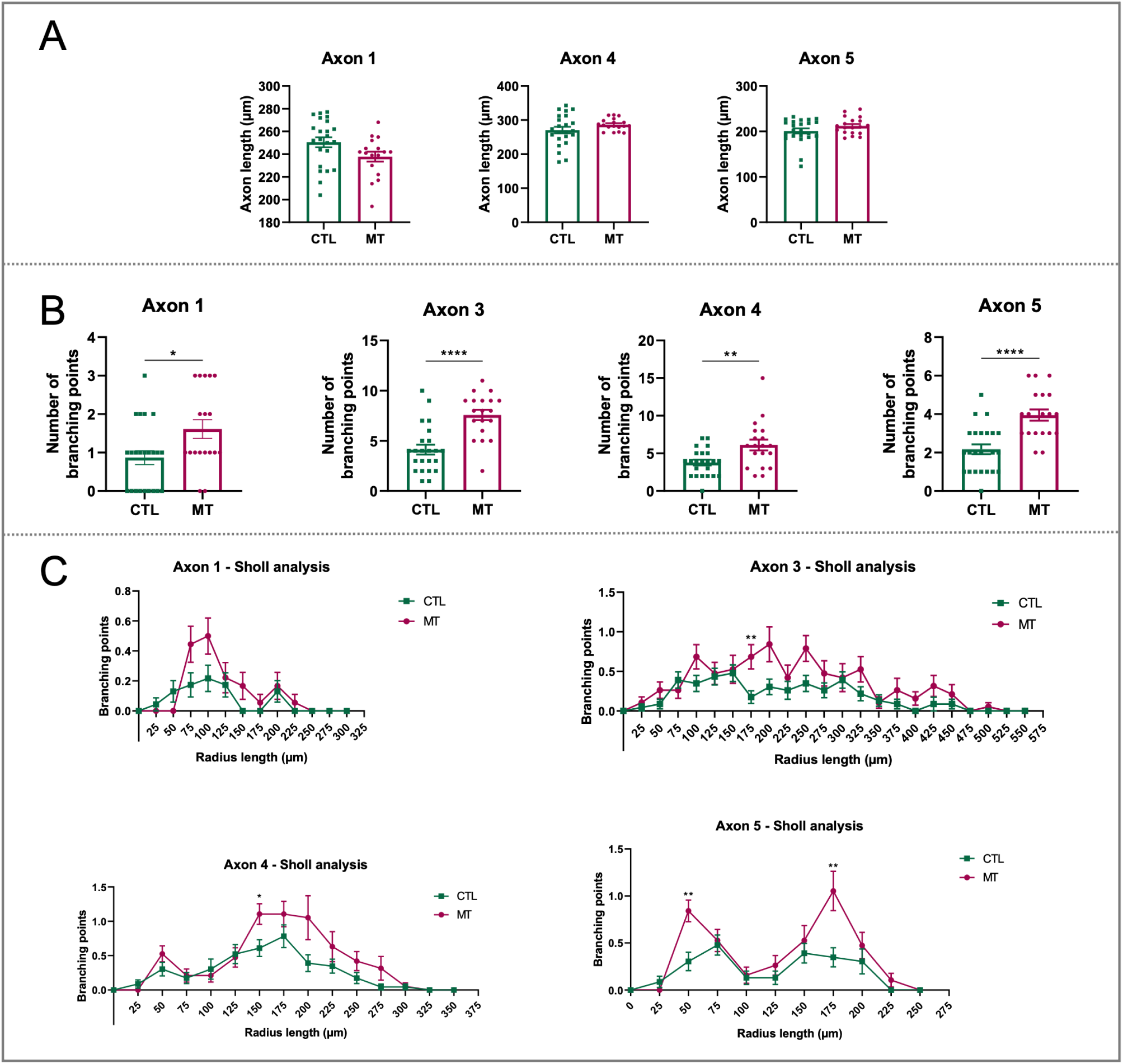
Complete tumour innervation analysis on D4. **A.** The graphs show the axon length on D4 of the axon number 1, 4 and 5 of CTL and MT larvae as indicated. The axon length is expressed in μm. **B.** The graphs represent the number of branching points on D4 of the axon number 1, 3, 4, and 5 of CTL and MT larvae as indicated. Differences among groups (**A-B**) were analysed by two-tailed unpaired Student’s t test: * P-value<0,05, ** P-value <0,01. **C**: Sholl profiles of axon number 1, 3, 4, and 5 on D4 of CTL and MT larvae. Differences among groups were analysed by Mann-Whitney test. Results are expressed as means ± SEM. n=18-23. CTL = larvae injected with PBS MT (melanoma transplant) = larvae transplanted with A375P GFP + cells.

**Sup. Fig. 4:**
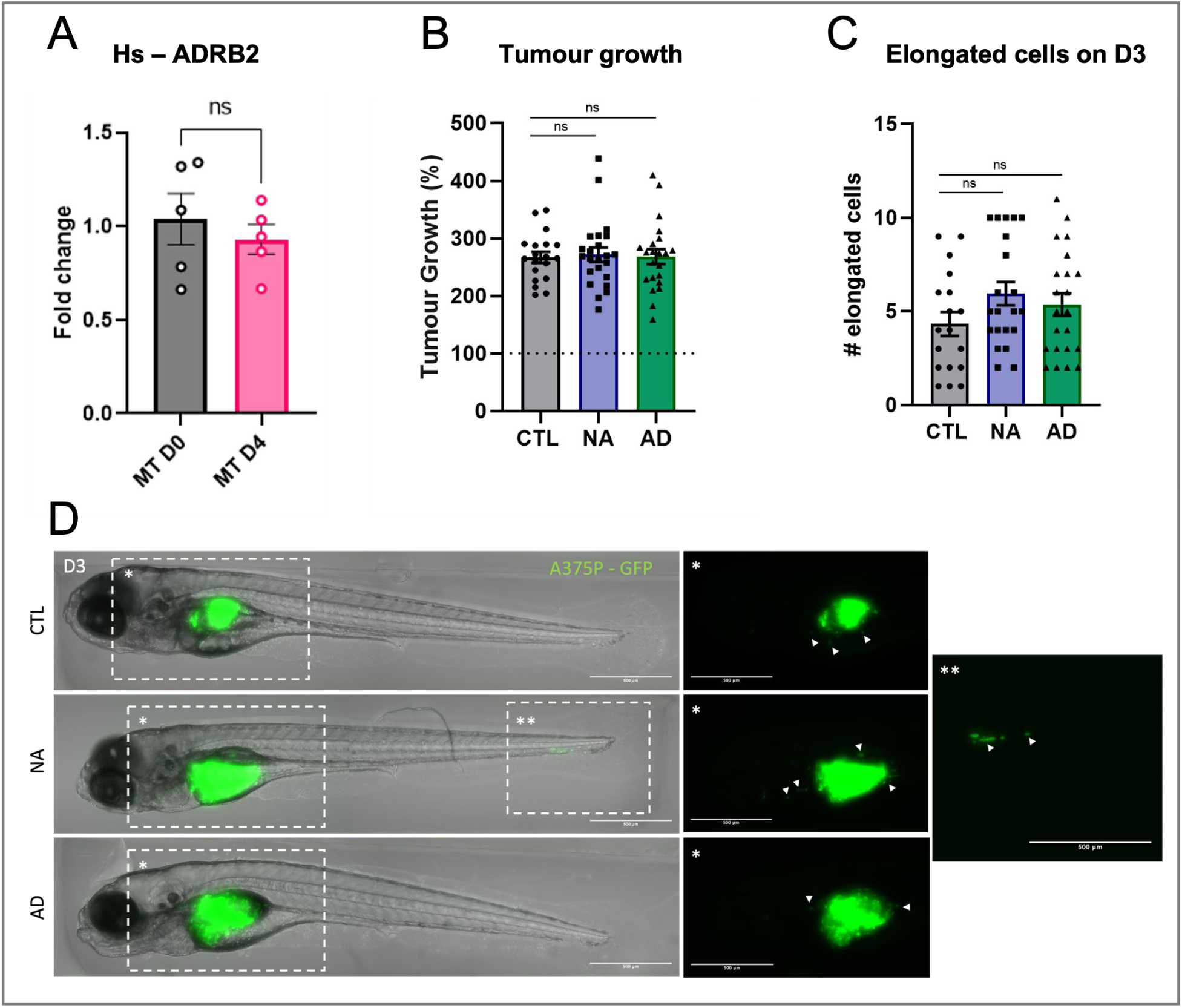
Role of catecholamines in xenograft melanoma model. **A.** Quantitative PCR analysis of the expression of human beta-adrenergic receptor 2 (*ADRB2*) in larvae transplanted with A375P cells on D0 and D4. Two-tailed unpaired Student’s t test was performed to evaluate the significance. n=5. **B.** Tumour growth analysis upon co-injection with catecholamines. The tumour area of individualized larvae was monitored from D0 to D3. The values are expressed as percentage of tumour growth, they are normalized on the tumour area of D0 for every separated condition. The tumour area is expressed in μm^2^. n= 19-23. **C.** The graph shows the quantification of elongated cells on D3. Elongated cells were manually counted in larvae for every condition on D3. n=18-24. Differences among treated conditions and control condition (**B-C**) were analysed by ordinary one-way ANOVA. Results are expressed as means ± SEM. **D.** Representative images of transplanted larvae with A375P cells (green) co-injected with PBS (CTL), 10 μM noradrenaline (NA) and 1 μM adrenaline (AD). First column: z-projection of full larvae. Dashed rectangles point area with detached cells that are zoomed on the following images. Second/third columns: zoomed z-projection of the selected areas where detached cells are present. Scale bar = 500 μm. CTL = larvae transplanted with A375P cells and co-injected with PBS. NA 10 μM = larvae transplanted with A375P cells and co-injected with 10 μM of noradrenaline. AD 1 μM = larvae transplanted with A375P cells and co-injected with 1 μM of adrenaline.

## Author contribution

Conceptualization: FL, LF, KK Investigation: FL

Writing – Original Draft: FL

Writing – Reviewing & Editing: FL, MCM, MAN, BSL, LF, KK,

Supervision: MAN, MCM, LF, KK,

Funding Acquisition: NEUcrest-ITN Project (EU’s Horizon 2020 grant No. 860635), AZELEAD

## References

1. Saginala, K., Barsouk, A., Aluru, J. S., Rawla, P. & Barsouk, A. Epidemiology of Melanoma. Med Sci (Basel) 9, 139–149 (2021).

2. Chapman, P. B. et al. Improved Survival with Vemurafenib in Melanoma with BRAF V600E Mutation. N Engl J Med 364, 2507 (2011).

3. Hodi, F. S. et al. Improved Survival with Ipilimumab in Patients with Metastatic Melanoma. N Engl J Med 363, 711 (2010).

4. Davis, L. E., Shalin, S. C. & Tackett, A. J. Current state of melanoma diagnosis and treatment. Cancer Biol Ther 20, 1366–1379 (2019).

5. Kozar, I., Margue, C., Rothengatter, S., Haan, C. & Kreis, S. Many ways to resistance: How melanoma cells evade targeted therapies. Biochimica et Biophysica Acta (BBA) - Reviews on Cancer 1871, 313–322 (2019).

6. Wang, H., Yang, L., Wang, D., Zhang, Q. & Zhang, L. Pro-tumor activities of macrophages in the progression of melanoma. Hum Vaccin Immunother 13, 1556–1562 (2017).

7. Mabeta, P. Paradigms of vascularization in melanoma: Clinical significance and potential for therapeutic targeting. Biomedicine & Pharmacotherapy 127, 110135 (2020).

8. Dror, S. et al. Melanoma miRNA trafficking controls tumour primary niche formation. Nat Cell Biol 18, 1006–1017 (2016).

9. Bapat, A. A., Hostetter, G., Von Hoff, D. D. & Han, H. Perineural invasion and associated pain in pancreatic cancer. Nat Rev Cancer 11, 695–707 (2011).

10. Guo, K. et al. Interaction of the sympathetic nerve with pancreatic cancer cells promotes perineural invasion through the activation of STAT3 signaling. Mol Cancer Ther 12, 264–273 (2013).

11. Demir, I. E., Friess, H. & Ceyhan, G. O. Neural plasticity in pancreatitis and pancreatic cancer. Nat Rev Gastroenterol Hepatol 12, 649–659 (2015).

12. Reavis, H. D., Chen, H. I. & Drapkin, R. Tumor Innervation: Cancer Has Some Nerve. Trends in Cancer vol. 6 1059–1067 Preprint at 10.1016/j.trecan.2020.07.005 (2020).

13. Shurin, M. R., Shurin, G. V., Zlotnikov, S. B. & Bunimovich, Y. L. The Neuroimmune Axis in the Tumor Microenvironment. The Journal of Immunology 204, 280–285 (2020).

14. Magnon, C. et al. Autonomic nerve development contributes to prostate cancer progression. Science (1979) 341, 1236361 (2013).

15. Magnon, C. Role of the autonomic nervous system in tumorigenesis and metastasis. Mol Cell Oncol 2, e975643 (2015).

16. Cole, S. W., Nagaraja, A. S., Lutgendorf, S. K., Green, P. A. & Sood, A. K. Sympathetic nervous system regulation of the tumour microenvironment. Nat Rev Cancer 15, 563–572 (2015).

17. Kuol, N., Stojanovska, L., Apostolopoulos, V. & Nurgali, K. Role of the nervous system in cancer metastasis. J Exp Clin Cancer Res 37, (2018).

18. Faulkner, S., Jobling, P., March, B., Jiang, C. C. & Hondermarck, H. Tumor neurobiology and the war of nerves in cancer. Cancer Discov 9, 702–710 (2019).

19. Prazeres, P. H. D. M. et al. Ablation of sensory nerves favours melanoma progression. J Cell Mol Med 24, 9574–9589 (2020).

20. Vats, K. et al. Sensory Nerves Impede the Formation of Tertiary Lymphoid Structures and Development of Protective Antimelanoma Immune Responses. Cancer Immunol Res 10, 1141–1154 (2022).

21. Balood, M. et al. Nociceptor neurons affect cancer immunosurveillance. Nature 611, 405 (2022).

22. Magnon, C. et al. Autonomic nerve development contributes to prostate cancer progression. Science (1979) 341, 1236361 (2013).

23. Tibensky, M. & Mravec, B. Role of the parasympathetic nervous system in cancer initiation and progression. Clinical and Translational Oncology 2020 23:4 23, 669–681 (2020).

24. Janik, M. E. et al. Diversified β-2-adrenergic Receptor Expression and Action in Melanoma Cells. Anticancer Res 37, 3025–3033 (2017).

25. Calvani, M. et al. Norepinephrine promotes tumor microenvironment reactivity through β3-adrenoreceptors during melanoma progression. Oncotarget 6, 4615–4632 (2015).

26. Shimizu, A. et al. Prognostic significance of β2-adrenergic receptor expression in malignant melanoma. Tumour Biol 37, 5971–5978 (2016).

27. Scheau, C. et al. Neuroendocrine Factors in Melanoma Pathogenesis. Cancers (Basel) 13, 2277 (2021).

28. De Giorgi, V., Geppetti, P., Lupi, C. & Benemei, S. The Role of β-Blockers in Melanoma. Journal of Neuroimmune Pharmacology 15, 17–26 (2020).

29. Konantz, M. et al. Zebrafish xenografts as a tool for in vivo studies on human cancer. Ann N Y Acad Sci 1266, 124–137 (2012).

30. Marines, J., Lorenzini, F., Kissa, K. & Fontenille, L. Modelling 3D Tumour Microenvironment In Vivo: A Tool to Predict Cancer Fate. Curr Issues Mol Biol 45, 9076–9083 (2023).

31. Kimmel, C. B., Ballard, W. W., Kimmel, S. R., Ullmann, B. & Schilling, T. F. Stages of embryonic development of the zebrafish. Developmental Dynamics 203, 253–310 (1995).

32. Hanahan, D. & Weinberg, R. A. Hallmarks of cancer: The next generation. Cell 144, 646–674 (2011).

33. Rébé, C. & Ghiringhelli, F. Interleukin-1β and Cancer. Cancers (Basel) 12, 1–31 (2020).

34. Yuan, A., Chen, J. J. W., Yao, P. L. & Yang, P. C. The role of interleukin-8 in cancer cells and microenvironment interaction. Front Biosci 10, 853–865 (2005).

35. Santoriello, C. et al. Kita Driven Expression of Oncogenic HRAS Leads to Early Onset and Highly Penetrant Melanoma in Zebrafish. PLoS One 5, e15170 (2010).

36. Peri, F. & Nüsslein-Volhard, C. Live Imaging of Neuronal Degradation by Microglia Reveals a Role for v0-ATPase a1 in Phagosomal Fusion In Vivo. Cell 133, 916–927 (2008).

37. Wang, W. et al. Nerves in the Tumor Microenvironment: Origin and Effects. Front Cell Dev Biol 8, 1630 (2020).

38. Rima, M. et al. Dynamic regulation of the cholinergic system in the spinal central nervous system. Sci Rep 10, 1–13 (2020).

39. Nikaido, M. et al. Early development of the enteric nervous system visualized by using a new transgenic zebrafish line harboring a regulatory region for choline acetyltransferase a (chata) gene. Gene Expression Patterns 28, 12–21 (2018).

40. Rocha, M., Singh, N., Ahsan, K., Beiriger, A. & Prince, V. E. Neural crest development: insights from the zebrafish. Developmental Dynamics 249, 88–111 (2020).

41. Kim, C. H. et al. Zebrafish elav/HuC homologue as a very early neuronal marker. Neurosci Lett 216, 109–112 (1996).

42. An, M., Luo, R. & Henion, P. D. Differentiation and maturation of zebrafish dorsal root and sympathetic ganglion neurons. Journal of Comparative Neurology 446, 267–275 (2002).

43. Pawolski, V. & Schmidt, M. H. H. Neuron-Glia Interaction in the Developing and Adult Enteric Nervous System. Cells 10, 1–20 (2020).

44. Espinosa-Medina, I. et al. Parasympathetic ganglia derive from Schwann cell precursors. Science (1979) 345, 87–90 (2014).

45. Stemple, D. L. & Anderson, D. J. Isolation of a stem cell for neurons and glia from the mammalian neural crest. Cell 71, 973–985 (1992).

46. Mancino, M., Ametller, E., Gascón, P. & Almendro, V. The neuronal influence on tumor progression. Biochimica et Biophysica Acta (BBA) - Reviews on Cancer 1816, 105–118 (2011).

47. Coelho, M. et al. β-Adrenergic modulation of cancer cell proliferation: available evidence and clinical perspectives. J Cancer Res Clin Oncol 143, 275–291 (2017).

48. Yang, E. V et al. Norepinephrine upregulates VEGF, IL-8, and IL-6 expression in human melanoma tumor cell lines: implications for stress-related enhancement of tumor progression. Brain Behav Immun 23, 267–275 (2009).

49. Moretti, S. et al. β-adrenoceptors are upregulated in human melanoma and their activation releases pro-tumorigenic cytokines and metalloproteases in melanoma cell lines. Lab Invest 93, 279–290 (2013).

50. Pedri, D., Karras, P., Landeloos, E., Marine, J. C. & Rambow, F. Epithelial-to-mesenchymal-like transition events in melanoma. FEBS J 289, 1352–1368 (2022).

51. Murayama, E. et al. Tracing hematopoietic precursor migration to successive hematopoietic organs during zebrafish development. Immunity 25, 963–975 (2006).

52. Chen, L. et al. A zebrafish xenograft model for studying human cancer stem cells in distant metastasis and therapy response. Methods Cell Biol 138, 471–496 (2017).

53. Astell, K. R. & Sieger, D. Zebrafish In Vivo Models of Cancer and Metastasis. Cold Spring Harb Perspect Med 10, 1–17 (2020).

54. Lois, C. & Alvarez-Buylla, A. Long-Distance Neuronal Migration in the Adult Mammalian Brain. Science (1979) 264, 1145–1148 (1994).

55. Kulkarni, S. et al. Adult enteric nervous system in health is maintained by a dynamic balance between neuronal apoptosis and neurogenesis. Proc Natl Acad Sci U S A 114, E3709–E3718 (2017).

56. Ayala, G. E. et al. Cancer-related axonogenesis and neurogenesis in prostate cancer. Clin Cancer Res 14, 7593–7603 (2008).

57. Mauffrey, P. et al. Progenitors from the central nervous system drive neurogenesis in cancer. Nature 569, 672–678 (2019).

58. Lu, R. et al. Neurons generated from carcinoma stem cells support cancer progression. Signal Transduct Target Ther 2, 1–10 (2017).

59. Zhang, D. et al. Stem cell and neurogenic gene-expression profiles link prostate basal cells to aggressive prostate cancer. Nat Commun 7, 1–15 (2016).

60. Hashemi, G., Dight, J., Khosrotehrani, K. & Sormani, L. Melanoma Tumour Vascularization and Tissue-Resident Endothelial Progenitor Cells. Cancers (Basel) 14, 4216 (2022).

61. James, J. M. & Mukouyama, Y. suke. Neuronal action on the developing blood vessel pattern. Semin Cell Dev Biol 22, 1019–1027 (2011).

62. Larrivée, B., Freitas, C., Suchting, S., Brunet, I. & Eichmann, A. Guidance of Vascular Development. Circ Res 104, 428–441 (2009).

63. Dai, H. et al. Enhanced survival in perineural invasion of pancreatic cancer: an in vitro approach. Hum Pathol 38, 299–307 (2007).

64. Ayala, G. E. et al. Growth and Survival Mechanisms Associated with Perineural Invasion in Prostate Cancer. Cancer Res 64, 6082–6090 (2004).

65. De Giorgi, V., Geppetti, P., Lupi, C. & Benemei, S. The Role of β-Blockers in Melanoma. J Neuroimmune Pharmacol 15, 17–26 (2020).

